# The G3BP stress-granule proteins reinforce the integrated-stress-response translation program

**DOI:** 10.1101/2024.10.04.616305

**Authors:** Jarrett Smith, David P. Bartel

## Abstract

When mammalian cells are exposed to stress, they coordinate the condensation of stress granules (SGs) through the action of proteins G3BP1 and G3BP2 (G3BPs) and, simultaneously, undergo a massive reduction in translation. Although SGs and G3BPs have been linked to this translation response, their overall impact has been unclear. Here, we investigate the question of how, and indeed whether, G3BPs and SGs shape the stress translation response. We find that SGs are enriched for mRNAs that are resistant to the stress-induced translation shutdown. Although the accurate recruitment of these stress-resistant mRNAs does require the context of stress, a combination of optogenetic tools and spike-normalized ribosome profiling demonstrates that G3BPs and SGs are necessary and sufficient to both help prioritize the translation of their enriched mRNAs and help suppress cytosolic translation. Together these results support a model in which G3BPs and SGs reinforce the stress translation program by prioritizing the translation of their resident mRNAs.

## INTRODUCTION

When exposed to stresses, such as oxidative stress, high temperature, and harmful chemicals, cells initiate the integrated stress response (ISR).^1–4^ The ISR integrates signals from different stress sensors to converge on phosphorylation of translation initiation factor eif2α,^1–5^ which causes reduced global translation, while allowing selective translation of stress-responsive mRNAs that help cope with the stress.^2,4,6–9^ During this dramatic alteration of the translation state, cytoplasmic puncta known as stress granules (SGs) form through the process of liquid-liquid phase separation.^1,6,10–15^ These granules are non-membrane-bound collections of RNA-binding proteins, mRNAs, and translation machinery.^1,11,16–19^ Although SGs contain a diverse proteome, their formation requires the RNA-binding proteins G3BP1 and G3BP2 (G3BPs).^12,16^ Both SGs and G3BPs have been hypothesized to play roles in translation,^16,20–23^ but their precise impacts are not well understood.

Because they are highly enriched for translation machinery and form concurrently with translation shutdown, SGs were originally hypothesized to be required for the dramatic translation shutdown observed during the ISR.^20,24,25^ However, G3BP-KO cell lines, which lack microscopically visible SGs, still undergo the eif2α-dependent translation reduction associated with the ISR.^16,26^

Because the formation of SGs is enhanced by polysome collapse, and SGs are enriched in small ribosomal subunits but not large ribosomal subunits, SGs are also proposed to simply be sites devoid of translation.^1,2,10,20,27,28^ However, single-molecule analyses show that translating mRNAs are not forbidden inside of SGs,^29^ and some report that translation inside of SGs might not be a rare event compared to translation in the cytosol.^30^

Despite their many connections to translation and the stress response, the overall impact that G3BPs and SGs have on translation during stress or, indeed, whether they have any impact is not fully understood.^10,31–33^ Previous attempts to determine the effects of G3BPs or SGs on translation have relied either on bulk measurements of translation with limited sensitivity and specificity, such as S^35^ labeling of nascent peptides, or have been restricted to observations of single mRNAs and reporters.^16,21,30^ Here, we use spike-normalized ribosome profiling and optogenetic methods to measure the impact of G3BPs and SGs on translation with or without stress. These results support a model in which G3BPs and SGs reinforce the ISR translation program by prioritizing the translation of their most enriched transcripts.

## RESULTS

### Ribo-spike Enables Observation of Absolute Translation Changes during the ISR

Global changes to translation such as those observed during the ISR have been challenging to observe by ribosome profiling which typically lacks information on the absolute differences between samples.^34^ RNA spike-ins typically used to quantify absolute differences in other high-throughput approaches are not appropriate for ribosome profiling, as the RNAse treatment used to create ribosome-protected fragments (RPFs) would destroy the unprotected spike molecules. Normalizing to endogenous mitochondrial RPFs can bypass this problem.^34,35^ However, this approach assumes that mitochondrial translation is unaffected by the experimental condition—an assumption that does not always hold, especially if cells are subjected to stress. To use ribosome profiling to measure absolute changes to TE occurring as a consequence of stress, we developed a ribosome profiling spike-in (ribo-spike) consisting of a defined amount of rabbit reticulocyte lysate translating orthogonal firefly luciferase mRNA, which could then be spiked into each sample (Fig 1A and Extended Data Fig. 1A).

**Figure 1.**
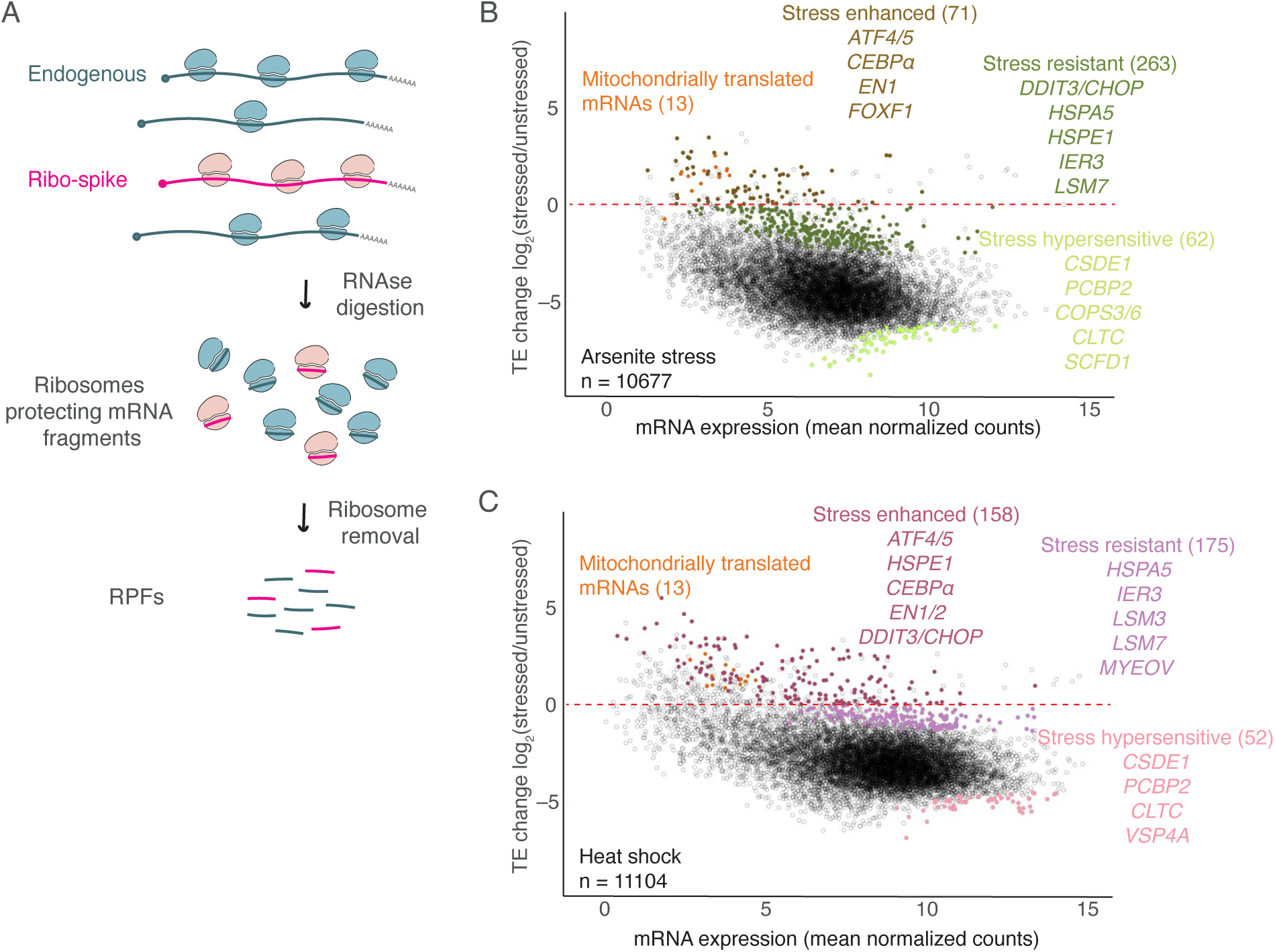
Ribo-spike enables observation of absolute translation changes during ISR. **(A)** Schematic representation of the ribo-spike. The ribo-spike consists of polysomes formed on an orthogonal mRNA sequence, in this case firefly luciferase mRNA (*fLuc*) translated in a rabbit reticulocyte lysate. A defined amount of the ribo-spike sample is added to each experimental sample, ultimately generating orthogonal RPFs that enable normalization between samples. (**B**) The ISR translation program induced upon arsenite stress. Plotted for each mRNA is the log2 fold-change (FC) in TE observed in HCT116 cells responding to 500 µM sodium arsenite (for 1 h) as a function of the mRNA expression. The red dashed line indicates unchanged TE, as determined by the ribo-spike. Points for stress-enhanced, stress-resistant, and stress-hypersensitive mRNAs are indicated by colors, with exemplar mRNAs listed for each category. Stress response categories were determined by DESeq2 (log2 FC > 1, adjusted *p* value < 0.05). Mitochondrially translated mRNAs are indicated in orange. *n* indicates number of unique mRNAs. **(C)** The ISR translation program induced upon heat shock (45°C for 25 min); otherwise, as in (A).

Using our ribo-spike, we then compared translation between unstressed cells and cells treated with 500 µM sodium arsenite for 1 h. This dose of sodium arsenite activates the ISR, leading to eif2α phosphorylation and substantial reduction in global translation.^16,20^ By normalizing to the spike-in TE across samples we were able to measure absolute changes in TE for mRNAs of >10,000 individual genes. Aggregating these measurements revealed a drastic (20-fold) reduction in global TE, consistent with previous bulk measurements (Fig 1B and Extended Data Fig. 1B,C).^16,36,37^ Interestingly, TEs of mitochondrial mRNAs increased in the presence of sodium arsenite (Fig 1B), validating our concern that they might not remain constant under stress.

Although translation was globally reduced during stress, some mRNAs retained or even enhanced their translation in response to stress. Among them were mRNAs of several stress-responsive genes previously reported to maintain their translation during the ISR, including *ATF4*, *ATF5*, *CEBP*α, and *DDIT3*/*CHOP* (Fig 1B). Many of these mRNAs contain upstream open reading frames (uORFs), which can help regulate their translation during stress.^38^ Indeed, regulation by uORFs is proposed to help shape the general translation response to stress.^2,4,7,8,39–41^ Supporting this proposal, mRNAs reported to contain one or more functional uORF were significantly better translated during the stress response (Extended Data Fig.1D).^42^ Moreover, while our manuscript was in review, another study published development of a similar ribo-spike and also applied it to show that uORF-containing mRNAs are better translated during stress.^41^

In yeast, newly transcribed mRNAs tend to be preferentially translated during stress.^43,44^ To assess this possibility in mammalian cells, we compared the translation change in response to stress to mRNA half-lives, reasoning that mRNAs with short half-lives would be predominantly composed of newly synthesized transcripts. Consistent with the observations made in yeast, we observed a moderate negative correlation between the translation change of an mRNA upon stress and its estimated half-life (Extended Data Fig. 1E; *R*_S_ = –0.22).^45^

At the other extreme, some mRNAs were especially sensitive to translation shutdown, including *CSDE1* and *PCBP2*, as reported previously.^40^ Interestingly, other hypersensitive mRNAs encoded proteins involved in energetically expensive processes, such as vesicle trafficking (*COPS3*, *COPS6*, *CLTC*, *SCFD1*), which are strongly down-regulated during stress (Fig 1B).^46^

To measure absolute TE changes in another stress context, we performed analogous experiments under 45°C heat shock (Fig 1C and Extended Data Fig. 1B,C). The results resembled those observed for arsenite stress (Fig 1C and Extended Data Fig. 1F,G). However,, the translation changes were not identical (Spearman *R*, *R*_S_ = 0.63), and many mRNAs were called as differentially regulated in one stress but not the other (Extended Data Fig. 1H–J), consistent with previous reports that the type and dosage of stress shape the ISR.^3^

We consider the combination of widespread translation shutdown, the retained or increased translation of select transcripts, and the hypersensitive downregulated translation of others to be the ISR translation program. These results illustrate how using ribo-spike enables accurate and precise monitoring of absolute translation changes between samples and emphasize the utility of a truly orthogonal spike-in for studies monitoring global changes to translation.

### G3BPs Reinforce the ISR Translation Program

Having developed a method to globally monitor absolute TE changes, we wanted to use it to determine the effects of G3BPs and SGs on the ISR. To do so, we needed to be able deplete cells of G3BPs and SGs. Therefore, we engineered HCT116 cells in which both endogenous G3BP1 and endogenous G3BP2 were homozygously tagged with RFP and an auxin-inducible degron (AID) allowing their rapid and efficient depletion (>90% within 3 h) upon addition of indole-3-acetic acid (IAA).^47,48^ By assessing another SG-marker protein, PABPC1 labeled with GFP, we confirmed that our depletion of G3BPs prior to stress imparted minimal effects on PABPC1 stability and largely prevented formation of SGs in response to both sodium arsenite and heat shock (Fig 2A–F).

**Figure 2.**
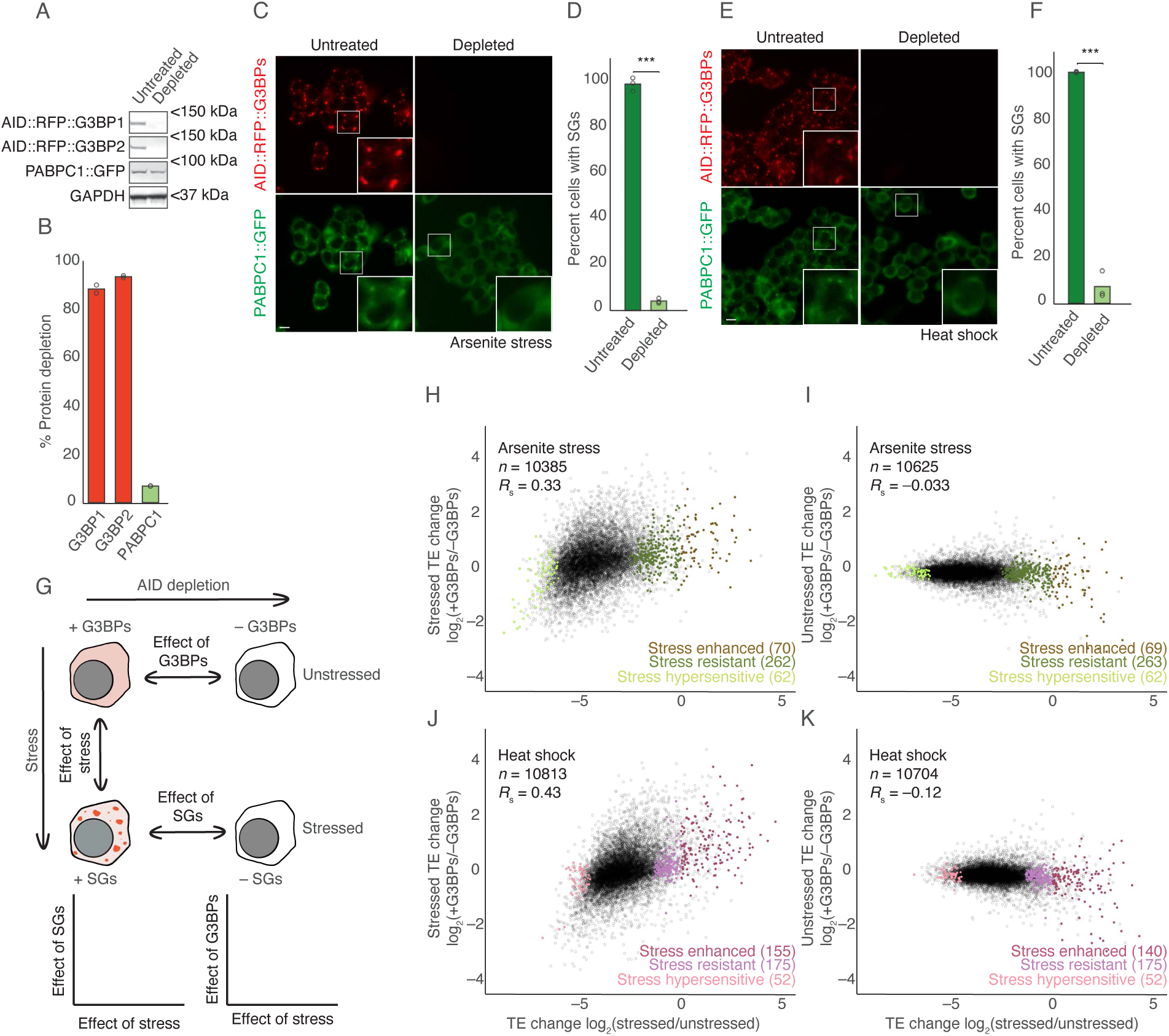
G3BPs reinforce the ISR translation program. **(A)** Depletion of endogenous G3BPs fused to AID. Immunoblots probed for the indicated proteins show specific depletion of AID fusion proteins after treatment with 500 µM IAA for 3 h. **(B)** Quantification of protein depletion in (A), normalizing to levels of GAPDH. Points show values for two biological replicates. **(C)** Prevention of SG formation during arsenite stress. HCT116 cells were either treated with 500 µM IAA to deplete G3BPs (right) or not treated (left) prior to arsenite stress (500 µM sodium arsenite for 1 h). Images show either G3BPs in red or SG marker protein PABPC1 in green (scale bar, 10 µm). **(D)** Quantification of SG formation in (C). Points show values for three biological replicates (***, *p* = 6.5 x 10^−5^; Welch’s two-sample, two-tailed, t-test). **(E, F)** Prevention of SG formation during heat shock (45°C for 25 min); (***, *p* = 0.001) otherwise, as in (C, D). **(G)** Schematic representation of G3BP-depletion experiment. Ribosome profiling was performed on stressed HCT116 cells to determine the effects of stress on translation, G3BP-depleted unstressed HCT116 cells to determine the effect of G3BPs on translation, and G3BP-depleted stressed HCT116 cells to determine the effects of G3BPs/SGs on translation. The effects of stress were then compared to the effects of G3BPs and SGs. **(H)** Relationship between G3BP-dependent translation during arsenite stress and the ISR translation program. The plots indicate the log2 FC in TE due to presence of G3BPs in cells treated with arsenite as a function of the log2 FC in TE due to arsenite. Points for stress-enhanced, stress-resistant, and stress-hypersensitive mRNAs are indicated by colors. *n* indicates number of unique mRNAs. **(I)** Relationship between G3BP-dependent translation in the absence of stress and the ISR translation program; otherwise, as in (H). **(J)** Relationship between G3BP-dependent translation during heat shock and the heat-shock ISR translation program; otherwise, as in (H). **(K)** Relationship between G3BP-dependent translation in the absence of stress and the ISR translation program; otherwise, as in (H).

Having generated these degron lines, we used ribosome profiling with the ribo-spike to compare translation in cells with and without the ability to form SGs (Fig 2G). Compared to G3BP-depleted cells, normal cells underwent a somewhat stronger inhibition of global translation during arsenite stress (1.4-fold, Extended Data Fig. 2A), consistent with reports that G3BPs are largely dispensable for the global reduction in translation caused by the ISR.^16^

We were intrigued to find that G3BPs and perhaps SGs did seem to contribute, if only modestly, to the massive reduction in translation that occurs during the ISR. To further examine this contribution, we compared the translation changes caused by presence of G3BPs to those caused by activation of the ISR. Here, we observed a clear positive correlation (*R*_S_ = 0.33; Fig 2H and Extended Data Fig. 2B). This implied that G3BPs, and perhaps SGs, tend to translationally upregulate the same mRNAs that the ISR upregulates and tend to translationally downregulate the same mRNAs that the ISR downregulates, thereby reinforcing the ISR translation program.

To determine whether these G3BP-dependent changes to TE were driven by effects on RPFs or RNA, we examined these two measurements separately. Interestingly, changes to RPFs and mRNA levels both contributed to the overall TE effects, with presence of G3BPs leading to more RPFs of ISR-enhanced transcripts despite a reduction in the levels of these transcripts (Extended Data Fig. 2C,D). Although these G3BP-dependent decreases in mRNA might represent an independent function of G3BPs, they are perhaps more parsimoniously explained as the translation-dependent destabilization of mRNA, which has been previously reported.^49–54^ Indeed, we also observed evidence of such mRNA stabilization occurring transcriptome-wide in response to arsenite stress (Extended Data Fig. 2E). However, the previously observed correlation between translation change upon stress and unstressed mRNA half-lives might complicate this interpretation (Extended Data Fig. 1E).

Given that the ISR translation programs differed between arsenite and heat-shock, we examined whether G3BPs also reinforced the heat shock ISR translation program and found similar results (Fig 2J, and Extended Data Fig. 2F–J). Furthermore, analyses of published measurements of G3BP-dependent translation in U2OS cells treated with the ER stressor thapsigargin, also yielded similar results (Extended Data Fig. 3A–E).^55^ Taken together, these results indicated that G3BPs, and perhaps SGs, reinforce the ISR translation program across multiple stresses and cell types.

We next asked whether the factors correlating with ISR translation—uORF presence and estimated mRNA half-lives—also correlated with G3BP-dependent translation during both arsenite stress and heat shock. However, we found no strong relationship between either of these factors and G3BP-dependent translation (Extended Data Fig. 4A – D).

To determine whether the G3BP-dependent translation was constitutive or stress-specific— potentially through formation of SGs, we performed analogous ribosome profiling experiments, comparing translation in cells with and without G3BPs, but this time in the absence of stress.

Interestingly, in unstressed cells, G3BP-dependent translation changes showed no positive correlation to the translation changes of arsenite or the heat-shock ISRs (*R*_S_ = –0.033 and –0.12 respectively; Fig 2I, K). Similar results were observed when analyzing published data from G3BP-KO U2OS cells (Extended Data Fig. 3B).^55^ Together, these results argued that G3BPs reinforce the ISR translation program in a stress-dependent manner, consistent with a model in which this is a function of G3BP through its nucleation of SGs.

Because G3BPs are required for SG formation, we could not say whether the effects we observed upon depleting G3BPs were a consequence of losing SGs or whether they were a consequence of losing some other stress-specific G3BP function. We sought to disambiguate G3BP and SG function by targeting CAPRIN1 or UBAP2L, two other proteins reported to be required for SG formation.^16,19,56,57^ However, no reduction in SG formation was detected upon AID depletion of either of these proteins (Extended Data Fig. 4E – M). These results concurred with findings that, in many cell types, G3BPs are uniquely required for SG formation, and illustrated the difficulties in disentangling the functions of SGs from those of their required G3BP components.^12^

### SGs are Enriched for mRNAs that are Favored by the ISR Translation Program

After difficulty using this genetic approach to define the role of SGs, we turned to other approaches. One attractive hypothesis was that mRNAs that are differentially regulated by the ISR translation program are also differentially localized to SGs, and that this localization to SGs determines the effect that G3BPs have on the translation of those mRNAs during stress.

To assess the relationships between regulation by the ISR, regulation by G3BPs, and SG localization, we purified stress granule cores from HCT116 cells treated with 500 µM sodium arsenite for one hour.^18,58^ Sequencing identified 488 transcripts enriched in SG cores and 453 that were depleted compared to total cytoplasm (Fig 3A, >2-fold change, adjusted *p* value <0.05). Overall, our SG enrichments observed in HCT116 cells resembled those previously reported from U2OS cells,^18^ including an enrichment for longer mRNAs and mRNAs that were poorly translated prior to stress. This indicated that we had successfully purified SG cores (Extended Data Fig. 5A – D).

**Figure 3.**
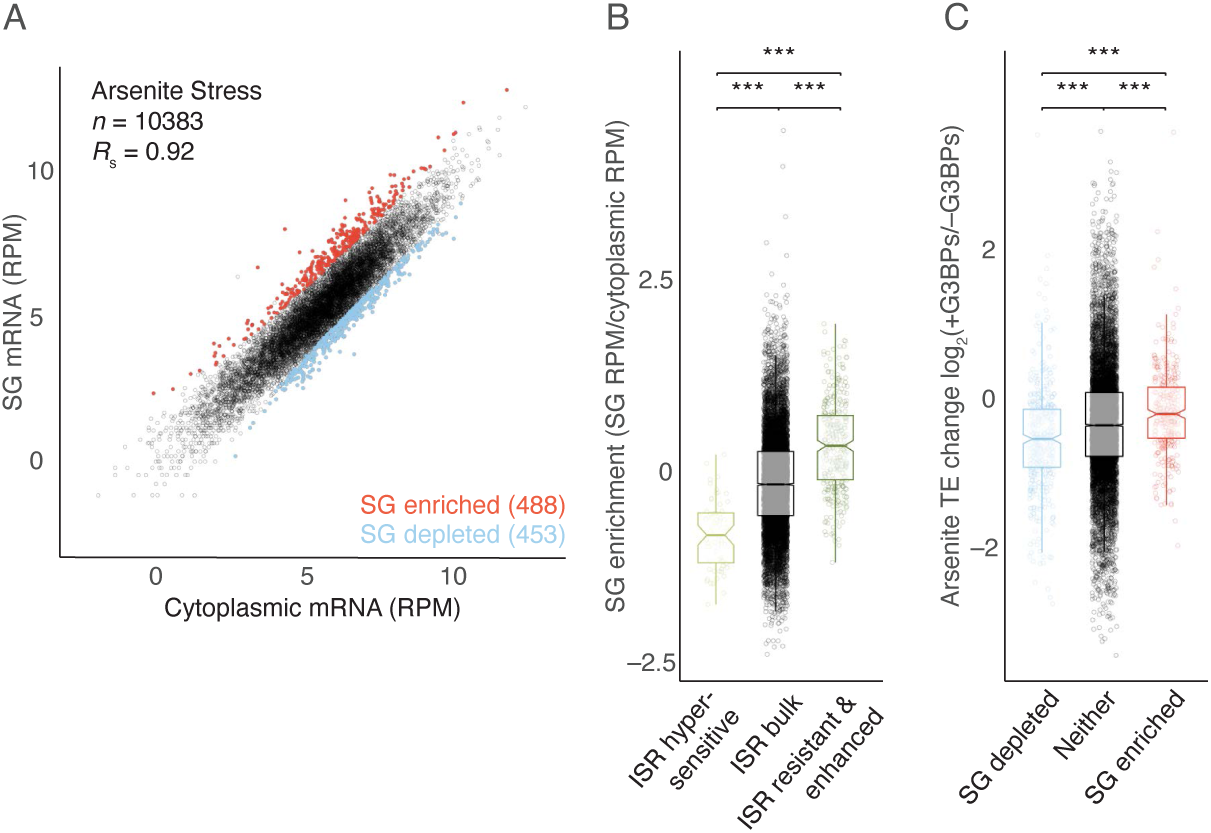
Translation of SG-enriched mRNAs is favored by G3BPs. **(A)** mRNAs enriched and depleted in SGs. Plotted for each mRNA is its abundance in the SG fraction as a function of its abundance in the cytoplasmic fraction. Points for SG-enriched and SG-depleted mRNAs, as determined by DESeq2 (log2 FC > 1, adjusted *p* value < 0.05), are indicated in red and blue, respectively, with their numbers in parentheses. *n* indicates total number of unique mRNAs. **(B)** Relationship between SG enrichment and ISR translation. Plotted are the SG enrichments of mRNAs in each of three ISR translation categories: stress-enhanced/resistant, bulk behavior, and stress-hypersensitive (line, median; notch, 95% confidence interval; box, quartiles; whiskers, 1.5 x IQR); left to right, ***, *p* = 7 x 10^−26^, 9 x 10^−56^, 6 x 10^−26^; Welch’s two-sample, two-tailed, t-test; n = 10,383 unique mRNAs). **(C)** Relationship between G3BP-dependent translation and SG enrichment. The plots indicate the log2 FC in TE due to presence of G3BPs in stressed cells for mRNAs in each of three SG-enrichment categories: SG-enriched, SG-depleted, and neither; left to right, ***, *p* = 2.6 x 10^−5^, 1.2 x 10^−15^, 5.6 x 10^−15^; otherwise as in (B). All stresses were 500 µM sodium arsenite for 1 h.

We next examined the relationship between ISR regulation and SG enrichment and found that mRNAs that retained or enhanced their translation during stress tended to be more enriched in SGs, whereas hypersensitive mRNAs tended to be depleted (Fig 3B). Furthermore, analyses of SG-enrichment in U2OS cells treated with sodium arsenite^18^, stress-dependent RNA-granule enrichment in NIH 3T3 cells treated with thapsigargin^59^, and stress-dependent G3BP proximity labeling in HEK293T cells treated with sodium arsenite^60^ yielded similar, albeit sometimes weaker, trends (Extended Data Fig. 5F – J). Together, these results supported a model in which mRNAs that are favored by the ISR translation response preferentially localize to SGs and suggest that this model might be generalizable across multiple stresses and cell types.

### SG-Enriched mRNAs are Favored by G3BPs During Stress

We next examined the relationship between G3BP regulation and SG enrichment in our HCT116 cells. Surprisingly, the presence of G3BPs tended to enhance TE of SG-enriched mRNAs during arsenite stress, whereas the presence of G3BPs tended to repress TE of mRNAs depleted from SGs during arsenite stress (Fig 3C). Moreover, repeating this analysis with SG enrichment data from arsenite-treated U2OS cells^18^, RNA-granule enrichment data from heat-shocked NIH 3T3 cells^59^, and G3BP proximity labeling data from arsenite-stressed HEK293T cells,^60^ all yielded analogous results (Extended Data Fig. 4K – M). Thus, in a variety of stresses and cell-types, the G3BPs appeared to favor the translation of SG-enriched mRNAs or disfavor the translation of SG-depleted mRNAs, suggesting a model in which those transcripts that are most strongly upregulated by the ISR translation program localize to SGs, leading to further upregulation by G3BPs.

### SG-like OptoGranules Drive Global Reduction in Translation without a Full ISR

Our global analyses showed that G3BPs, while not required to establish the ISR translation program, do reinforce it. To determine whether SG formation was indeed sufficient to drive these translation changes in the absence of exogenous stress, we set out to form ectopic SGs in the absence of exogenous stress. Overexpressing reported SG-nucleating proteins UBAP2L and G3BP1^61,62^ failed to induce authentic SGs in a majority of cells. UBAP2L-induced puncta lacked the canonical SG marker eIF4G, and, while puncta induced by G3BP1 overexpression did resemble authentic SGs, they formed in only ∼20% of cells (Extended Data Fig. 6A – D).

To more robustly form ectopic SGs, we adapted a published CRY2 optogenetic system.^63^ We replaced the N-terminal NTF2L dimerization domain of G3BP1 with the blue light-dependent cryptochrome 2 photolyase homology region oligomerization domain (CRY2), tagged with GFP (GFP::CRY2::G3BP_ΔN_) (Fig 4A). Even in the absence of blue light, doxycycline-induced expression of this construct caused formation of ectopic puncta in ∼20% of cells, similar to overexpressing GFP::G3BP1. However, exposing these cells to 488 nm blue light enhanced puncta formation, such that they were observed in ∼80% of cells within 3 h. Consistent with previous findings,^63^ our light-induced granules were also reversible (disassembling to basal levels within 2 h of blue light removal), positive for canonical SG markers eIF4G and CAPRIN1, and exhibited photobleaching dynamics resembling those of canonical SGs (Fig 4A,B, Extended Data Fig. 6E – G). Together, these results showed that our OptoGranules (OGs) were dynamic, reversible, SG-like condensates formed in the absence of exogenous stress.

**Figure 4.**
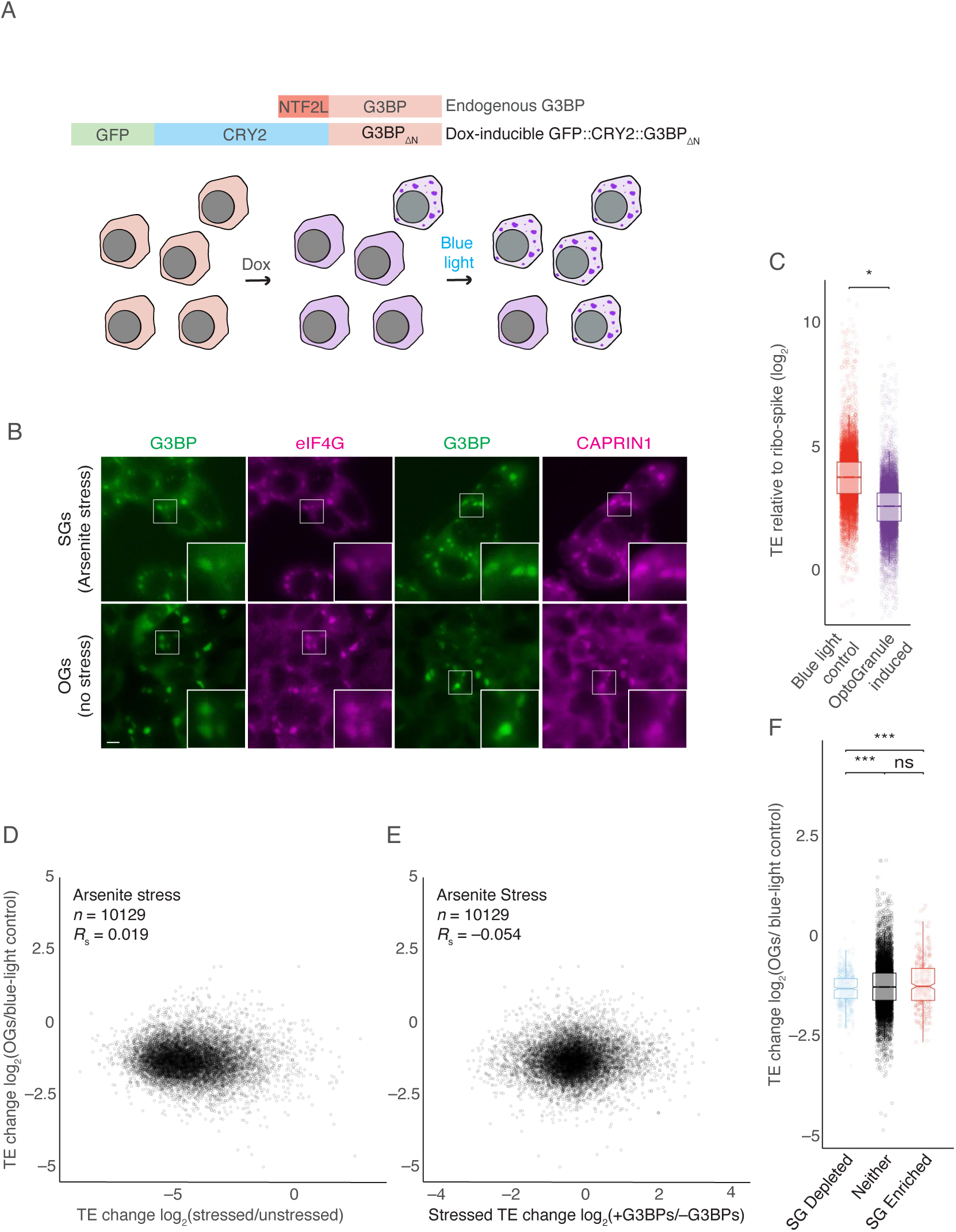
SG-like OGs Can Drive a Global Reduction in Translation. **(A)** Schematic representation of the OG system. HCT116 cells expressing endogenous G3BP containing an NTF2L dimerization domain were edited to also express doxycycline (dox)-inducible ectopic G3BP in which the NTF2L domain had been replaced with a fusion of GFP and the light-inducible CRY2 dimerization domain (GFP::CRY2::G3BPΔN). After doxycycline induction, a minority of cells formed ectopic granules. After exposure to blue light (488 nm), most cells formed ectopic granules. **(B)** SG marker localization to OGs. HCT116 cells expressing wild-type GFP::G3BP1 were either stressed with 500 µM sodium arsenite for 1 h (top) or treated with 1 µM doxycycline to induce expression of GFP::CRY2::G3BPΔN and exposed to blue light for 3 h (bottom). Fluorescent or immunostained proteins are as indicated (scale bars, 10 µm). White boxes highlight insets that are expanded on the right. n = 3 replicates with similar results. (**C**) Comparison of TE in OG-induced cells to that in blue-light control cells. Plotted are the TE distributions in HCT116 cells in which GFP::CRY2::G3BPΔN was expressed and exposed to blue light to form OGs (right) and in control cells expressing only wild-type G3BP exposed to blue light, in which no OGs were formed (left) (line, median; notch, 95% confidence interval; box, quartiles; whiskers, 1.5 x IQR; n = 10,129 unique mRNAs). Significance was determined using ribo-spike values as in Figure S1B (*, *p* = 0.026; Welch’s two-sample, two-tailed, t-test). **(D)** Comparison of OG and ISR translation programs. Plotted is TE change caused by induction of OGs as a function of TE change caused by arsenite. *n* indicates total number of unique mRNAs. **(E)** Comparison of OG and G3BP-dependent translation programs. Plotted is TE change caused by OGs as a function of TE change caused by presence of G3BPs in arsenite-treated cells; otherwise, as in (C). **(F)** Comparison of OG translation program and SG enrichment. The plots indicate the log2 FC in TE caused by induction of OGs for mRNAs in each of three SG enrichment categories: SG-enriched, SG-depleted, and neither (line, median; notch, 95% confidence interval; box, quartiles; whiskers, 1.5 x IQR; n = 10,129 unique mRNAs); ns, not significant; left to right, ***, *p* = 6.9 x 10^−9^, 2.3 x 10^−5^, 0.067; Welch’s two-sample, two-tailed, t-test).

If SG-like condensation alone was sufficient to drive an ISR-like response, then we would predict that induction of OGs would yield the following results. First, cells with OGs would undergo a global reduction in translation. Second, these OG-induced translation changes would positively correlate with ISR-induced translation changes. Third, OG-induced translation changes would also positively correlate with G3BP-dependent translation during stress. Fourth, transcripts enriched in SGs would be resistant to any translation shutdown caused by the formation of these SG-like condensates.

To test these predictions, we induced OGs in HCT116 cells, and used ribosome profiling with our ribo-spike to compare them to cells exposed to blue light but lacking GFP::CRY2::G3BP_ΔN_, and therefore lacking OGs. We observed a modest global TE reduction in OG-induced cells when compared to blue-light controls (Fig 4C). This modest reduction (1.6-fold) agreed with our earlier result that G3BPs were responsible for a small fraction of the ISR translation shutdown (1.4-fold for arsenite stress) (Extended Data Fig. 2A), and was consistent with the first of our four predictions. However, analyses of the TE results failed to confirm any of the remaining three predictions. OG translation changes did not correlate with either the ISR translation program or G3BP-dependent translation changes in stressed cells (Fig 4D,E, *R*_S_ = 0.019 and – 0.054, respectively), and mRNAs that were enriched in SGs did not perform any better than those that were neither enriched nor depleted (Fig 4F). Considered together, these results argued that condensation of the SG-like OGs, while sufficient to drive a modest reduction in global translation, was not sufficient to specify the ISR translation program.

### OptoGranules Require Cellular Stress to Recruit an SG Transcriptome

One caveat of our previous analysis was the assumption that spontaneously formed OGs contained the same transcriptome as SGs and could therefore regulate those transcripts in the same manner. This assumption aligned with evidence that G3BP was the central scaffold of the SGs complex protein–RNA interaction network and was further supported by the observation that OGs were positive for many canonical SG markers (Extended Data Fig. 6E).^12–14,63^ Interestingly, this assumption was also consistent with the observation that the established determinants of the SG transcriptome (transcript length and TE prior to stress) are parameters that are independent of stress.^18^ However, the SG transcriptome is also thought to be largely determined by the transcriptome RNA-binding landscape (the landscape created by proteins bound to the transcriptome).^10,12,64,65^ Stress can dramatically remodel this RNA-binding landscape, shifting the RNA content of other biological condensates, such as P-bodies and P granules.^66,67^ Thus, SGs formed in the presence of stress and OGs formed in the absence of stress might recruit different sets of mRNAs, despite their general compositional, morphological, and biophysical similarities.

To test this hypothesis, we purified OGs formed in the presence of stress. To do this, we integrated the doxycycline-inducible GFP::CRY2::G3BP_ΔN_ construct into cells expressing endogenously edited AID::RFP::G3BPs and doxycycline-inducible OsTIR1, the E3 ligase responsible for the turnover of the AID. Upon treatment with doxycycline and IAA, these cells degraded their endogenous G3BPs and replaced them with GFP::CRY2::G3BP_ΔN_, rendering them incapable of forming SGs in the absence of blue light, even when stressed. However, when these cells were stressed and exposed to blue light, they readily formed granules (Fig 5A,B). We refer to these OGs formed in the presence of stress as StressOptoGranules (SOGs).

**Figure 5.**
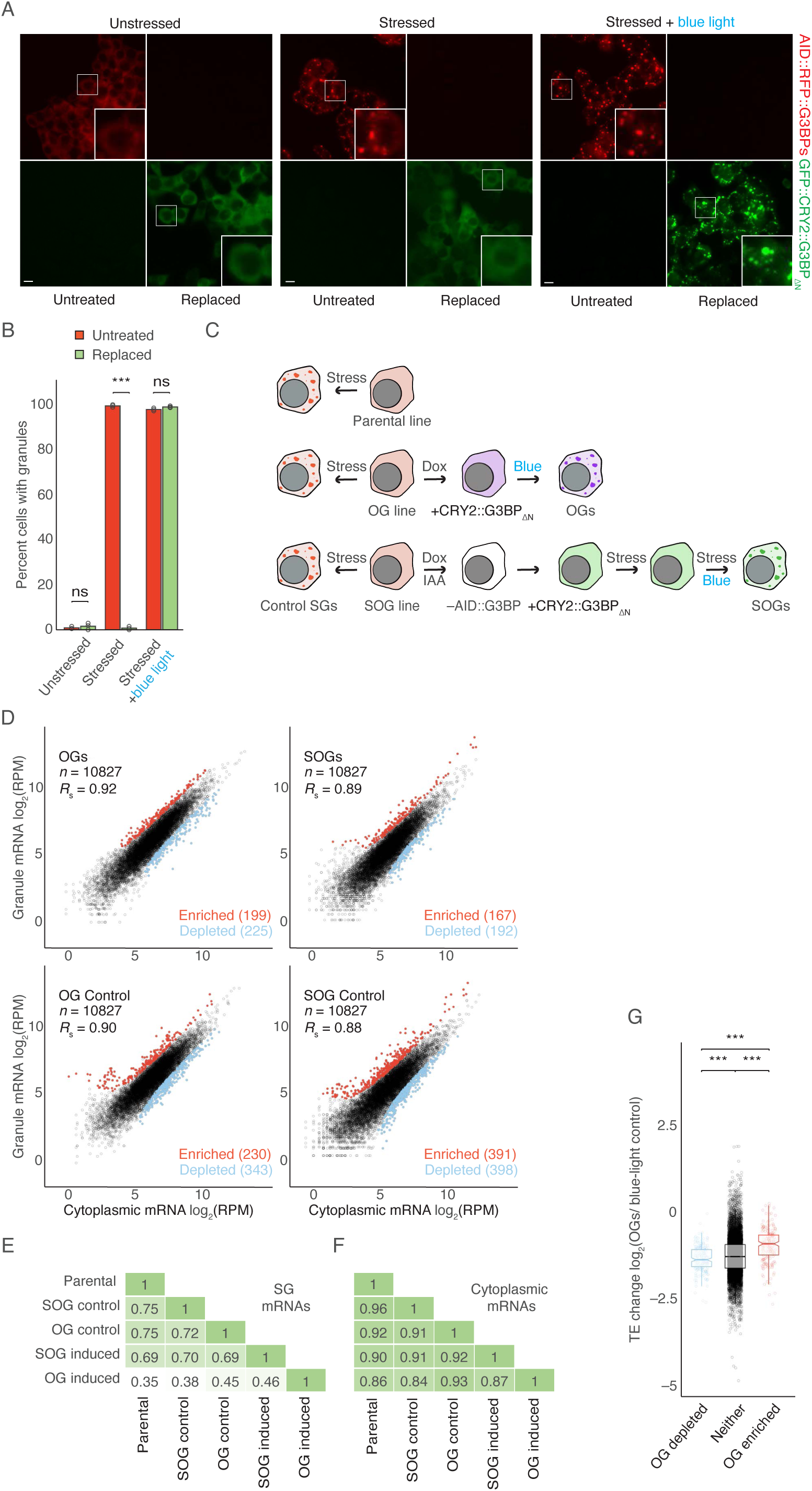
OGs require stress to recruit the SG transcriptome. **(A)** Replacement of G3BP with GFP::CRY2::G3BPΔN to induce SOGs. In the left panel, HCT116 cells expressing endogenously tagged AID::RFP::G3BPs were either untreated (left) or treated with 1 µM doxycycline and 500 µM IAA for 18 h (right), leading to turnover of endogenous G3BP shown in red and expression of GFP::CRY2::G3BPΔN shown in green. In the middle panel, cells were treated as in left panel, but were subsequently stressed with 500 µM sodium arsenite for 1 h in the absence of blue light. In the right panel, cells were treated as in the middle panel, but were also exposed to blue light, beginning 30 min after start of the arsenite treatment (scale bars, 10 µm). White boxes highlight insets that are expanded on the right. **(B)** Quantification of granule formation in (A). Error bars report standard deviation for three biological replicates (left to right, ns, *p* = 0.48; ***, *p* = 5.9 x 10^−9^; ns, p = 0.071; Welch’s two-sample, two-tailed, t-test). **(C)** Schematic representation of granule purification experiment. Three cell lines were used: the parental HCT116 line expressing endogenous G3BPs, an OG HCT116 line expressing endogenous G3BPs and doxycycline-inducible GFP::CRY2::G3BPΔN, and an SOG line expressing endogenous G3BP fused to AID and RFP (RFP::AID::G3BP), doxycycline-inducible OsTIR1, and doxycycline-inducible GFP::CRY2::G3BPΔN. Control SGs were purified from each cell line by treating them with 500 µM sodium arsenite for 1 h. Additionally, OGs were purified from the OG line treated with doxycycline for 18 h, followed by 3 h of blue light, and SOGs were purified from the SOG line treated with both doxycycline and IAA for 18 h followed by exposure to arsenite stress for 30 min in the absence of blue light and then an additional 30 min of stress in the presence of blue light. **(D)** mRNAs enriched and depleted in the indicated types of granules; otherwise, as in Fig. 3A. **(E)** Comparisons of granule enrichments. The heatmap depicts the pairwise correlations (*R*S values) observed between the enrichments of granules from (Fig. 3A and 5D). **(F)** Comparisons of cytoplasmic transcriptomes of cells used to determine granule enrichments (E). **(G)** Comparison of OG-dependent translation and OG enrichment. Plotted are distributions of TE changes caused by induction of OGs for mRNAs in each of three OG enrichment categories; left to right, ***, *p* = 9.5 x 10^−9^, 1.5 x 10^−17^, 9.5 x 10^−11^; otherwise, as in Fig 4F.

We then purified and sequenced the transcripts associated with both OGs and SOGs. We compared transcripts across OGs, SOGs, and control SGs from each cell line including our original SG enrichment data acquired from the parental line (Fig 5C,D). Control SGs correlated strongly with each other (*R*_S_ = 0.72–0.75). OGs formed in the absence of stress, correlated substantially less with control SGs (*R*_S_ = 0.35–0.45), indicating a distinct transcriptome (Fig 5E, Extended Data Fig. 7A). Interestingly, we found that transcripts enriched in SOGs strongly resembled those enriched in SG controls (*R*_S_ = 0.69–0.70), indicating that the difference between SG and OG transcriptomes was due to lack of cellular stress, as opposed to some intrinsic bias of OG formation (Fig 5E, Extended Data Fig. 7A). These differences in granule composition did not appear to be driven by differences in gene expression, as the cytoplasmic mRNA levels correlated quite well between these groups (*R*_S_ = 0.84–0.96) (Fig 5F, Extended Data Fig. 7B). Taken together, our results indicated that establishing the SG transcriptome requires not only the interaction network established by G3BPs but also the RNA-binding landscape established during cellular stress.

These OG enrichment data also allowed us to confirm that the transcripts enriched in OGs did indeed better retain their translation (Fig 5G). Taken together, our OG ribosome-profiling and purification results supported a model in which SGs are sufficient to drive a modest decrease in global translation while prioritizing the translation of their enriched mRNAs. However, SGs required stress to induce an ISR-like translation program, presumably because the sorting of SG mRNAs is dictated by the cellular RNA-binding landscape, which is drastically remodeled upon activation of the ISR.

### Tethering to SGs Imparts Resistance to ISR Translation Shutdown

Our findings suggested that localization to an SG grants an mRNA prioritized translation during the ISR. To test this model, we examined whether tethering a reporter to SGs influenced its translation. The reporter encoded nanoluciferase (nLuc) fused to the *E. coli* dihydrofolate reductase (ecDHFR) destabilizing domain, which causes rapid turnover of its protein fusions, ensuring reporting on recently translated nLuc.^68^ The 3’UTR of the reporter mRNA included an array of 24 bacteriophage MS2 hairpins, which bind MS2 coat protein (MCP), thereby providing a means to tether the reporter to the SG (Fig 6A).^30,69^

**Figure 6.**
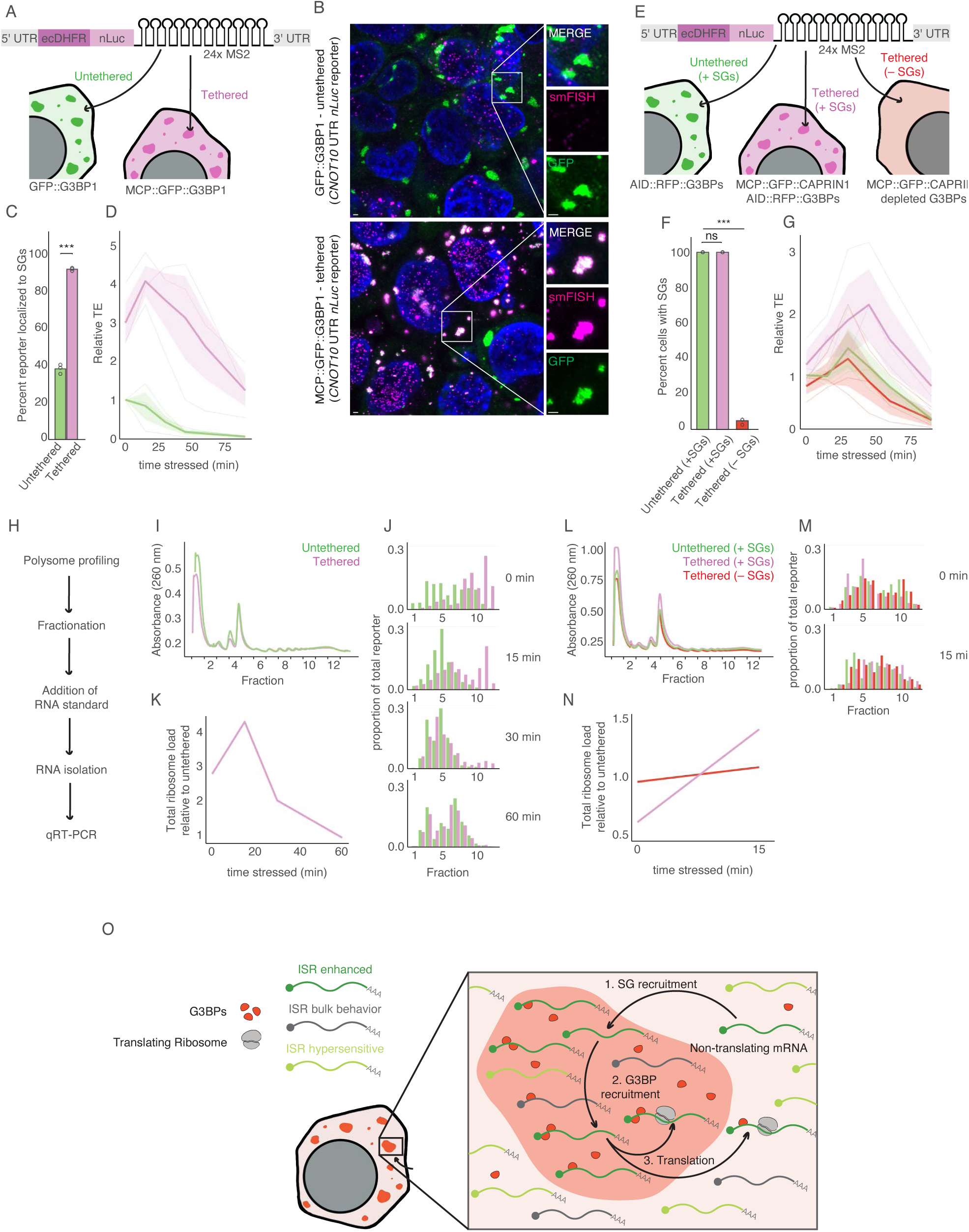
Tethering to SGs imparts resistance to ISR translation shutdown. **(A)** Schematic representation of G3BP1 tethering. **(B)** Tethering a luciferase reporter to SGs via G3BP1. HCT116 cells expressing a NanoLuc reporter bearing 5’ and 3’ UTRs from *CNOT10* and endogenous G3BP1 tagged with either GFP or MCP::GFP were stressed with 500 µM sodium arsenite for 90 min. Images show G3BP1 in green and smFISH of NanoLuc reporter molecules in magenta (scale bars, 1 µm). White boxes highlight insets on the right. n = 3 replicates with similar results. **(C)** G3BP1 tethering efficiency. Percent of SG-localized reporter molecules in either untethered cells (green) or tethered cells (pink) are shown. Points show values for biological replicates (***, *p* = 3.6 x 10^−6^; Welch’s two-sample, two-tailed, t-test; n = 3 biological replicates). **(D)** G3BP1-tethered ISR translation. Plotted is the TE of the reporter in untethered cells (green) or tethered cells (pink) during a time course of arsenite stress (500 µM sodium arsenite). Bold line shows average values from at least three biological replicates shown as thin lines. Light-colored ribbons report standard error. TE is reported relative to that of untethered, unstressed cells from the matched biological replicate. **(E)** Schematic representation of CAPRIN1 tethering. **(F)** Quantification of SG formation in cells treated as depicted in (E). Error bars report standard deviation for three biological replicates (ns, *p* = 1; ***, *p* = 5.9 x 10^−8^; Welch’s two-sample, two-tailed, t-test). **(G)** CAPRIN1-tethered ISR translation. Plotted is the relative TE of the reporter in untethered cells (green), tethered cells with SGs (+SGs, pink), and tethered cells without SGs (–SGs, red); otherwise as in (D). **(H)** Schematic representation of polysome profiling experiment. We performed polysome fractionation on cells treated with 500 µM sodium arsenite, added an in vitro transcribed RNA standard to each fraction, isolated total RNA, and performed RT-qPCR to measure the abundance of the reporter relative to the standard. **(I)** Polysome profile analysis of MCP::GFP::G3BP1 expressing cells. Plotted are the polysome traces produced by the workflow depicted in (H). X-axis shows 12 equal-volume fractions that were taken with boundary points between fractions marked as tics. **(J)** Reporter distributions across G3BP1-tethered polysome profile fractions. Plotted are the proportions of total reporter mRNA in each polysome fraction as measured in (H,I) by RT-qPCR; otherwise as in (I). **(K)** Relative ribosome loading of G3BP1-tethered reporter. Plotted is the total ribosome load as determine by polysome profiling of the G3BP1-tethered reporter relative to its untethered counterpart. **(L)** Polysome profile analysis of MCP::GFP::CAPRIN1 expressing cells. Plotted are the polysome traces for untethered cells expressing endogenous G3BPs (green), tethered cells expressing MCP::GFP::CAPRIN1 along with endogenous G3BPs (pink), and tethered cells expressing MCP::GFP::CAPRIN1 with endogenous G3BPs depleted (red); otherwise as in (I). **(M)** Reporter distributions across CAPRIN1-tethered polysome profile fractions. Plotted are the proportions of total reporter mRNA in each polysome fraction of conditions shown in (L). Plots are shown for stress timepoints of 0 and 15 min; otherwise as in (J). **(N)** Relative ribosome loading of CAPRIN1-tethered reporter. Plotted is the total ribosome load as determine by polysome profiling of the CAPRIN1-tethered reporter in the presence of G3BPs (pink) and in their absence (red) relative to their untethered counterpart (green), before and during arsenite stress; otherwise as in (K). **(O)** A Plausible Model of SG Translation Control. 1) ISR-enhanced transcripts are preferentially recruited to SGs between cycles of translation, when they lack ribosomes. 2) As a consequence of being recruited to SGs, these transcripts are licensed for translation, possibly through the recruitment of G3BPs. 3) After this licensing, this model is agnostic to whether the transcript diffuses out to the cytosol, or remains inside the SG to be translated. Once these transcripts complete their translation, they would be available to, once again, be preferentially recruited to (or retained in) the granule.

G3BPs strongly partition into SGs^16,62,70^; almost half of our endogenous G3BP1 fusion protein (45.9%) localized to SGs after 90 min of arsenite stress (Extended Data Fig. 8A,B). Considering this high partitioning coefficient, we chose an MCP fusion to endogenous G3BP1 as our SG-tethering protein. Our reporter was stably integrated into HCT116 cells in which all endogenous G3BP1 alleles were tagged with either GFP (GFP::G3BP1) or MCP and GFP (MCP::GFP::G3BP1) (Fig 6A). We also explored the λN–BoxB aptamer system,^71,72^ but found that it perturbed G3BP localization and SGs (Extended Data Fig. 8C).

To assess the effects of SG tethering on an mRNA not normally recruited to SGs, we created a reporter using the 5’ and 3’ UTR of *CNOT10*, an mRNA that was not enriched in SGs in either our data or published datasets,^18^ and was translationally repressed both by the ISR and by G3BPs during stress (Table S1). Single-molecule fluorescence in situ hybridization (smFISH) with probes targeting the *nLuc* sequence indicated that the reporter was recruited to SGs at a basal level of 38% in cells expressing only GFP::G3BP (Fig 6B,C). In cells expressing MCP::GFP::G3BP, the reporter was more robustly recruited to SGs (92%) (Fig 6B,C), a higher fraction than that observed for 99% of endogenous mRNAs.^18^

Next, we measured how the translation of this reporter responded to arsenite stress. Cells were treated with 500 µM sodium arsenite for timepoints ranging from 0 to 90 min, and protein production was monitored by measuring luciferase signal. Reporter mRNA levels were then monitored by RT-qPCR to calculate a TE for each timepoint. For untethered reporter, we observed a 95% reduction in reporter TE over the 90 min period of stress (Fig 6D). This aligned with the 20-fold average decrease in TE observed by ribosome profiling, indicating that our assay accurately reported on stress-induced translation changes.

We then examined what happens to the translation of this reporter when tethered to MCP::GFP::G3BP1. Strikingly, even when tethered to G3BP1 in unstressed cells, our reporter was more efficiently translated, with an average 3-fold higher TE. This observation was consistent with previous reports that G3BPs can act as translational regulators, even outside the context of stress (Fig 6D).^16,21–23,73,74^ When stressed, we saw that our G3BP1-tethering continued to enhance the translation of our reporter. Interestingly, this translation enhancement peaked early in stress, as translation both in tethered and untethered cells diminished at later timepoints. However, even at our latest timepoints, our tethered reporter was translated significantly better its untethered counterpart (Fig 6D).

To test whether these effects on translation were specific to this reporter, we created two additional reporters using 5’ and 3’ UTRs from the genes *DDIT4L* and *RB1*, which represented a range of endogenous translational responses to stress and to G3BPs (Table S1). The results for both of these reporters resembled those observed for *CNOT10*, with the tethered reporter showing enhanced SG recruitment and translation across all stressed timepoints, peaking early in stress and then decreasing at later timepoints (Extended Data Fig. 8E – G).

We next wanted to determine whether, as in our ribosome profiling experiment, changes to mRNA levels contributed to these changes in translation efficiency. Examining protein and RNA separately, we saw that, although the rate of luciferase production was modestly higher for some tethered reporters, as evidenced by the slower rate of luciferase disappearance upon stress (Extended Data Fig. 8H,I), the untethered reporter mRNAs levels consistently increased across all reporter constructs during the stress treatment, by an average of 5.2-fold, whereas G3BP1-tethered mRNA levels remained steady or slightly decreased, with an average change of 0.9-fold (Extended Data Fig. 8J,K). This pattern was consistent with a model in which translation destabilizes the reporter mRNA, the stress-dependent translation shutdown stabilizes it, and recruitment to SGs, in rescuing translation, destabilizes the reporter yet again.^49,50,53,54^

To test this model, we treated cells expressing our untethered *CNOT10* reporter with two different translation inhibitors – cycloheximide (CHX) and puromycin (puro), and monitored protein and mRNA levels. As expected, both drugs inhibited translation by 98% over a 4 h time course (Extended Data Fig. 8L,M). Both drugs also caused dramatic stabilization of reporter mRNA (CHX by 10-fold; puro by 22-fold) (Extended Data Fig. 8N). These results indicated that our reporters are subject to translation-dependent destabilization, and are consistent with a model in which the translation-shutdown during stress contributes to the stabilization of our reporters.

To further distinguish between SG and G3BP function, we examined whether the effect of tethering was specific to G3BPs. To do this, tethered our *CNOT10* reporter to SGs using another SG protein, CAPRIN1. CAPRIN1-mediated recruitment to SGs was robust, resembling that observed with G3BP tethering (Extended Data Fig. 9A,B). To measure the translation effects of this alternative tethering strategy, we integrated both the reporter and doxycycline-inducible, MCP tagged CAPRIN1 into our AID::G3BP cell line. With this combination, we could assess the effects of tethering the reporter to SGs via CAPRIN1 and also deplete G3BPs to control for any effects that did not require formation of SGs (Fig 6E).

We first expressed the reporter in untreated cells without the induction of any MCP-tagged protein (Fig 6F, Extended Data Fig. 9C). In this condition, our reporter underwent a ∼75% reduction in translation efficiency, similar to our previous experiments (Fig 6G). We then induced MCP-tagged CAPRIN1 (MCP::GFP::CAPRIN1) without depleting the AID-tagged G3BPs by treating cells with both 1 µg/ml doxycycline and 200 µM auxinole, a small-molecule inhibitor of OsTIR1 activity (Fig 6F, Extended Data Fig. 9C). Under these conditions, the CAPRIN1-tethered reporter was more efficiently translated following arsenite treatment, indicating that tethering effects do not require tethering via G3BP (Fig 6G). However, these benefits to translation were abolished when cells expressing MCP::GFP::CAPRIN1 were depleted of G3BPs by treating cells with both 1 µg/ml doxycycline and 500 µM IAA (Fig 6F,G, Extended Data Fig. 9C), indicating that this effect of tethering required the presence of G3BPs and perhaps SGs.

As with our G3BP1-tethering experiments, we then replicated these CAPRIN1-tethering experiments using both the *DDIT4L* and *RB1* reporters. We observed results similar to those obtained with *CNOT10* at the TE, protein, and RNA levels, consistent with the idea that the reporter was better translated only when recruited to SGs and thus underwent translation dependent destabilization (Extended Data Fig. 9D – I). Interestingly, the average effect size observed with CAPRIN1 tethering was smaller than that observed with G3BP1 tethering, and, unlike our G3BP1-tethered reporters, CAPRIN1-tethered reporters showed only modestly improved translation in unstressed cells (Extended Data Fig. 9D,E). This difference may suggest that, while heavily G3BP-bound mRNAs are subject to enhanced translation, even in unstressed conditions, other mRNAs only benefit once they are recruited to the SG and brought into proximity with G3BPs.

To further measure the effect of SG association on translation by an orthogonal method less likely to be affected by mRNA stability or the kinetics of luciferase production and degradation, we monitored the ribosome association of our *CNOT10* reporter using polysome profiling RT-qPCR (Fig 6H). As expected, untethered reporter mRNA shifted gradually from heavy polysomes to lighter polysomes, and finally to monosomes and free RNA fractions over a 60 min arsenite treatment time course. However, G3BP-tethered reporter mRNA was associated with heavier polysome fractions compared to its untethered counterpart, even in unstressed cells, where SGs were not formed. Additionally, this preferential association peaked early in stress and diminished by the 60-min timepoint, when SGs were still fully formed but translation had largely shut down (Fig 6J,K). Both the preferential polysome association of our G3BP-tethered reporter in the absence of SGs, and the collapse in this preference at later time points suggested that this shift was not due to the inclusion of the reporter in heavy-sedimenting granules, but due to more efficient loading into actual ribosomes. These results indicated that tethering to G3BP enhanced reporter translation in both unstressed and stressed conditions, and were consistent with the results of our luciferase assays.

To further evaluate the results of our luciferase assays, we performed polysome profiling of the *CNOT10* reporter in CAPRIN1-tethered cells under unstressed and early-stress (15 min) conditions. Consistent with results of our luciferase assay, CAPRIN1 tethering enhanced ribosome association only during stress and in a G3BP-dependent manner (Fig 6L – N). The G3BP dependence of this polysome shift further supported the idea that it was not due simply to CAPRIN1 protein sedimenting farther into the gradient. Collectively, the results of these tethering experiments indicated that tethering the 3’UTR of an mRNA to either G3BP1 or CAPRIN1 increased recruitment to SGs and imparted preferential translation during the ISR— consistent with a model in which localization to SGs grants an mRNA prioritized translation during stress.

## DISCUSSION

SGs were once thought to be required for the global repression of translation observed during the ISR.^1,11,20,24,25^ Later studies hypothesized that SGs might have no function at all and are incidental byproducts of increased RNA availability due to translation repression.^32,33^ Here, we propose a model in which SGs are neither required for the ISR nor functionless condensates, but instead measurably enhance and reinforce the ISR translation program, thereby leading to a widespread and statistically significant, albeit relatively modest, impact on global translation.

This widespread yet subtle reinforcement of the ISR is consistent with several observations of SG biology. SGs recruit a wide array of mRNAs and proteins and yet contain only ∼10% of the cytoplasmic mRNAs and protein molecules.^10,17,18,59^ This implies that they would be more likely to subtly tune widespread gene expression than to have dramatic impacts. Indeed, both our ribosome profiling and reporter experiments demonstrate that, in addition to translational effects, SGs modestly regulate mRNA levels. Whether this additional example of SGs imparting subtle, global, tuning effects on gene expression was indirectly caused by translation-dependent destabilization of mRNAs or illustrates an independent function of SGs will be an interesting question for further investigation.

Consistent with the idea that SGs impart global tuning effects, a recent study investigating the impact of SGs during viral infection reports that SGs play a role in tuning the innate immune response to viral infection.^57^ However, instead of amplifying the ISR translation response, here SGs are reported to dampen the ISR transcriptional response.^57^ This difference in results might be explained by differences in the nature of the stressors or composition of the SGs.^19,37^ Compared to SGs formed by arsenite, heat shock, or ER stress, the granules formed in response to viral infection tend to be smaller, only weakly recruit PABPC1 and polyadenylated mRNAs, do not require the phosphorylation of eIF2α for their formation, and can include RNAse L bodies, which share many of the same markers with SGs and also form in response to viral infection.^57,75,76^ Moreover, the response to viral infection might differ between cells or viruses, as this negative tuning role has been absent in some other reports on viral stress.^77^ Thus, although our results seem likely to be generalizable across canonical SGs that require eIF2α phosphorylation, granules formed under viral infection appear to have distinct effects.

In presenting a model in which SGs reinforce the ISR translation program, we propose that SGs help to promote the translation of a subset of RNAs. Several observations have led to the hypothesis that SGs play strictly repressive roles in translation. However, none of these observations actually preclude SGs from promoting translation. Although SGs are nucleated by translationally silent mRNAs, implying that the mRNAs localized to them are non-translating, those mRNAs also dynamically exchange with the cytosol and could therefore transit out of the SG after their translation was initiated, maintaining enrichment for translationally silent mRNAs inside the SGs.^25,29,30,36,78^. Indeed, translation initiation itself might even expel the mRNA from the SG, as the presence of even a single ribosome has been reported to prevent SG localization.^79^ Similarly, although SGs are enriched for 40S but not 60S ribosomal subunits, implying that 60S subunits are not stoichiometrically available to assemble translating 80S ribosomes, 60S subunits are not depleted from SGs and appear to be equally available to form translating ribosomes as in the cytosol.^20,28^ Indeed, a recent single-molecule study reports that translation of an ATF4 reporter inside the SG is neither impossible nor rare.^30^ Thus, the prior observations, although originally interpreted as evidence of SGs being incompatible with translation, are also consistent with the possibility that they could be sites of privileged translation for a subset of mRNAs.

Although a model of mRNAs being preferentially translated while inside the SG is perhaps the simplest interpretation of our results, it is not the only interpretation. Alternatively, mRNAs could be licensed for translation while transiting through an SG. In this scenario, preinitiation complexes (PICs) or translational activators like UBAP2L, which are known to be enriched in SGs, could preferentially interact with mRNAs transiting through the SG.^1,10,20,28,56,80^ Then, having acquired this license, the mRNAs could go on to be translated independent of their localization inside or outside of the SG (Fig 6O). Such a model, in which licensed mRNAs are translated inside or outside the SG, could reconcile the previous observation that mRNAs devoid of ribosomes are preferentially recruited to SGs with our observation that mRNAs that are translationally enhanced during the ISR are enriched in SGs, further enhancing their translation. Such a model could also explain why a single-molecule reporter based on the *ATF4* mRNA is no more efficiently translated in the SG than in the cytosol.^30^ This being said, we found that *ATF4* mRNA was neither enriched in SGs nor influenced by the presence of SGs, indicating that other transcripts might better represent mRNAs influenced by SGs.

The difficulty of distinguishing the function of SGs from the function of the SG-nucleating G3BPs must be acknowledged. One approach to inferring SG function is to consider shared phenotypes of depleting proteins reported to be required for SG formation. However, many of these proteins are thought to also form complexes with each other, independent of SGs, obscuring the interpretation of shared phenotypes.^19,32,33^ Moreover, we and others found that the requirement for reported non-G3BP nucleators appears to be stress or cell-type specific, which limits the utility of depleting these factors.

Although we have not conclusively distinguished between these models, five lines of reasoning support the proposal that the ISR translation program is reinforced by the formation of SGs rather than by only G3BPs. First, both our ribosome profiling experiments and our tethering experiments demonstrated stress-specific effects. Although G3BP1 tethering enhanced translation in unstressed cells, this might reflect the effect of our 24xMS2/MCP system artificially recruiting many copies of G3BP simultaneously, and thereby simulating the SG environment. Second, although our SG purifications relied on G3BP IPs, these were IPs on SG-enriched fractions depleted of cytosolic G3BP.^17,18,58,81^ Thus, our data should reflect the SG transcriptome and not simply the G3BP–RNA interactome. Furthermore, we identified similar trends in SG-enrichment data that do not require IP of G3BPs.^59^ Third, the process of forming OptoGranules, which simply manipulates G3BPs condensation, was sufficient to trigger a translation response. Fourth, tethering reporter mRNAs to SGs, whether by G3BP1 or CAPRIN1, was sufficient to impart resistance to stress-induced translation shut down. Although G3BPs and CAPRIN1 are binding partners^16^, this result indicates that this effect does not require direct recruitment via G3BPs.

Lastly, in distinguishing between SG- and G3BP-centric models, we should not dismiss the possibility that both G3BPs and SGs contribute to this translation reinforcement. G3BPs have been observed to regulate translation and exhibit exceptionally high partitioning into SGs.^16,21–23,71,73,74^ Indeed, we observed almost half of G3BP1 localizing to SGs and that tethering to the protein enhanced translation, even in unstressed cells. If G3BPs license translation, perhaps recruitment of mRNAs to SGs is a molecular strategy to increase their interaction with G3BPs, reinforcing their translation during stress (Fig 6O). Although these considerations do not conclusively distinguish between SG and G3BP function, they do support the proposal that SGs are a functional condensate responsible for reinforcing the ISR translation program.

## Methods

### Cell culture

All cells were cultured at 37°C with 5% CO_2_ using McCoy’s 5A medium supplemented with 10% fetal bovine serum (FBS) and 2mM L-glutamine. HCT116 (CCL-247 from ATCC) were obtained from the Young lab.

### Cell-line construction

#### Cas9 mediated editing

Fusion proteins were introduced into the endogenous locus of genes using the Cas9 genome editing system. The 5’ and 3’ homology arms (each 200 – 700 bp) flanking an insertion site were either amplified from genomic DNA of the parental cell line or from gblock gene fragments (IDT) cloned into a rescue template plasmid, flanking the fusion protein sequence. This plasmid, along with pX330 plasmid expressing a Cas9 guide RNA, was reverse transfected into the parental cell line using lipofectamine 3000 transfection reagent in a 6-well dish at 100% confluency. After 48 h, these cells of each well were expanded to a 10 cm dish and cultured for an additional 24 h. Cells with strong signal from the fusion protein fluorescence were single-cell sorted into 96-well plates using flow cytometry. Colonies derived from single cells were grown, expanded, and screened by PCR genotyping and western blotting. Unless otherwise stated, all endogenously edited cell lines were edited homozygously.

#### PiggyBac editing

Exogenous genes were introduced into cells using the PiggyBac transposon system. The exogenous gene was either amplified from genomic DNA of the parental cell line or from gblock gene fragments (IDT) and cloned into a plasmid expressing the PiggyBac inverted-repeat sequences along with a selection marker for either hygromycin, puromycin, or blasticidin. This plasmid, along with the super PiggyBac transposase expression vector (System Biosciences #PB210PA-1), was reverse transfected into the parental cell line using lipofectamine 3000 transfection reagent (Thermo #L3000015) in a 6-well dish at 100% confluency. After 48 h, cells of each well were expanded to a 10 cm dish and successfully edited cells were selected for using the appropriate antibiotic. For cases in which a fluorescent protein had been transposed into the genome, cells expressing the protein at the appropriate level were single-cell sorted into 96-well plates by flow cytometry. Colonies derived from single cells were grown, expanded, and screened by either microscopy, western blotting, or both.

### Ribo-spike production

#### In vitro transcription

Ribo-spike mRNA was generated using mMessage mMachine kit (Thermo # AM1344) following the standard T7 protocol. DNA transcription template was generated by PCR of a plasmid template encoding the firefly luciferase coding region (*fLuc*) preceded by the T7 promotor and followed by an encoded poly(A) tail of 30 nt length. The resulting PCR product was purified using phenol–chloroform extraction followed by ethanol precipitation and was size selected and purified from an agarose gel using the “freeze and squeeze” method in which the gel slice was frozen at –80°C for 30 min, followed by centrifugation at 21,000*g* for 5 min.^82^ The eluted DNA was then concentrated by an additional ethanol precipitation and added to the mMessage mMachine in vitro transcription reaction, incubated for at 37°C for 2 h, and then treated with TURBO DNAse for 15 min at 37°C. RNA from the in vitro transcription reaction was recovered by passing the reaction over a tris-buffered micro bio-spin p30 gel column (Bio-Rad #7326223) centrifuging at 1000*g* for 4 min. The resulting flowthrough was then purified by phenol–chloroform extraction and ethanol precipitation, and then resuspended in water. RNA was examined by running an aliquot on a 4% urea–polyacrylamide denaturing gel and staining with Sybr Gold (Thermo #S11494).

#### In vitro translation

The *fLuc* mRNA was in vitro translated using a rabbit reticulocyte lysate system (Promega #L4151) following the standard non-nuclease treated protocol, with the exception of using a shorter incubation time to maximize ribosome occupancy. The reaction was assembled by combining 0.06 µM *fLuc* mRNA with reticulocyte lysate, 0.02 mM complete amino acid mixture, 0.8 units/µl RNAsein ribonuclease inhibitor, 0.01 M creatine phosphate, 0.05 mg/ml creatine phosphokinase, 2 mM DTT, 0.05 mg/ml tRNA, 80 mM potassium acetate, and 0.5 mM magnesium acetate in 23 117 µl reactions. The reactions were incubated at 30°C for 15 min, cooled on ice and treated with 0.1 mg/ml cycloheximide to block ribosome translocation. Reactions were then combined, mixed, aliquoted, snap frozen in liquid nitrogen, and stored at – 80°C.

### IAA-induced protein depletion

HCT116 cells were engineered using the PiggyBac system to express multiple copies of doxycycline-inducible *OsTIR1*. These cells were then also engineered using Cas9 to expressed AID-tagged fusion proteins edited at their endogenous loci. Cells expressing fusion proteins were first doxycycline-induced for 4 h and were then treated with 500 µM IAA (Gold Bio #I-110-25) to induce rapid (< 3 h) depletion of the AID-fusion protein. Depletion was confirmed by microscopy, western blot, or both. When confirming by western, as was the case with G3BP1, G3BP2, and CAPRIN1 proteins, antibodies raised directly against the target proteins were used to enable detection of untagged alleles, if present. Where indicated, non-depleted controls were treated with 200 µM auxinole (Aobious #AOB8812) to minimize background protein degradation.^83^ Unless otherwise stated, all other non-depleted control cells were treated only with ethanol, the solvent used for IAA. Previous work has shown that addition of IAA alone does not affect translation.^84^

### Stress conditions

For oxidative stress, unless otherwise stated, cells were treated with media containing 500 µM sodium arsenite (Sigma #93289) for 1 h. For heat shock, media was separately heated to 45°C. Media was removed from cells previously grown in a 37°C, rapidly replaced with 45°C media, and then cells were quickly placed in a 45°C incubator for 25 min.

### Doxycycline induction

For all doxycycline inductions, cells were treated with 1 µg/ml doxycycline (Clonetech Takara Bio #631311) for indicated amount of time.

### Tethered reporter TE experiments

To measure changes in TE for the nanoluciferase reporters (NanoLuc), cells expressing a reporter containing selected 5’ and 3’ UTRs, NanoLuc fused to an ecDHFR domain and 24x MS2 stem loops, were washed with cold PBS, lysed directly in their dish in buffer containing 10 mM Tris HCl pH 7.5, 5 mM MgCl2, 100 mM KCl, 1% triton, 1x cOmplete mini tablet (EDTA free), and 0.02 U/µl SUPERase-In RNase inhibitor, and depleted of nuclei by centrifuging at 1300*g* for 10 min at 4°C. Nuclear-depleted supernatant was then transferred to a new tube and flash frozen using liquid nitrogen and stored at –80°C. Cells were then assayed for total protein using bicinchoninic acid, luciferase production, and reporter mRNA using RT-quantitative PCR (RT-qPCR) comparing to the *GAPDH* mRNA, which did not change in abundance during treatment with sodium arsenite (Extended Data Fig. 8C). Translation efficiency was then calculated according to the following equation:

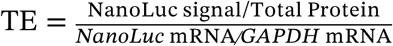

In G3BP-tethering experiments, the reporter was integrated into in two different HCT116-derived cell lines: a line expressing endogenous G3BP1 tagged with GFP (GFP::G3BP1), and a line expressing endogenous G3BP1 tagged with the MS2 coat protein (MCP) fused to GFP (MCP::GFP::G3BP1)(Fig 6A). In CAPRIN1-tethering experiments, the reporter was expressed in an HCT116-derived cell line under three different conditions: untethered, in which the cell line was untreated and expressed endogenous G3BPs tagged with AID and RFP (AID::RFP::G3BPs); tethered (+SGs), in which the cell line was treated with 1 µM doxycycline and 200 µM auxinole for 10 h, leading to retention of G3BPs and expression of MCP::GFP::CAPRIN1; and tethered (– SGs), in which the cell line was treated with 1 µM doxycycline and 500 µM IAA for 10 h, leading to turnover of endogenous G3BPs, which prevented SG formation, and expression of MCP::GFP::CAPRIN1 (Fig 6E).

#### Luciferase Assays

Luciferase assays were conducted using the standard protocol of the Luciferase Reporter Assay System (Promega #E1910). Cell lysates were analyzed in 96-well microplate format at room temperature using a GloMax microplate reader and luminometer device (Promega #GM3000). Each sample was run in technical duplicate.

#### RT-qPCR

RNA was extracted using TRI reagent (sigma #<u>93289</u>), precipitated using ethanol and linear acrylamide (thermo #J67830-XF) and resuspended in water. RT reactions were performed using a Quantitect RT kit following kit protocols. Genomic DNA was first eliminated using 7x gDNA wipeout buffer and incubating at 42°C for 2–5 min. Reactions were then placed on ice, RT reagents were added, and then RT was performed at 42°C for 30 min. RT enzyme was heat inactivated by incubating at 95°C for 3 min. cDNA product was then diluted 1:2 in water and stored at –20°C. qPCR was then performed using the PowerUP SYBR Green qPCR master mix and primers targeting either the *NanoLuc* or *GAPDH* sequences. Each sample was run in technical triplicate PCR using a QuantStudio 6 Pro RT-PCR System (Thermo).

### Blue-light LED treatment

Cells were exposed to blue light for indicated amounts of time using a 12 inch square LED Grow Light System with 225 x 14W Blue LEDs (HQRP #884667106091218) mounted inside of their cell culture incubator.

### Fluorescence recovery after photobleaching

Cells were bleached using a blue light (488 nm) laser, followed by green channel imaging every second for 3 min. Imaging was performed using an Andor Revolution Spinning Disk Confocal with FRAPPA system. FRAP Analysis was performed using Fiji (ImageJ).

### Immunofluoresence and live imaging

Cells were grown in plastic Nunc Cell Culture Treated Multidishes (Thermo #140675). For live imaging, cells were aspirated of their cell growth media and replaced with PBS. For immunofluorescence, cells were fixed using 3.7% formaldehyde in PBS at room temperature for 15 min, washed with PBS for 5 min three times, blocked using normal goat serum diluted in PBS with 0.3% triton for at least 1 h, then treated with primary antibody diluted in 1% normal goat serum, 0.3% triton, and PBS. Primary antibody treatments were performed overnight at 4°C. Cells were then washed in PBS for 5 min three times and treated with secondary antibody using same dilution buffer for 1–2 h at room temperature. Cells were washed a final time with PBS and mounted using mounting media containing DAPI (VECTASHIELD #H-2000-10). Imaging was performed on a Nikon Ti automated inverted microscope with incubation enclosure. Images were analyzed using Fiji (Image J).

### smFISH

For smFISH, cells were plated in glass bottom, 10 mm, 24 well plates (MatTek Corporation #P24G-1.5-10-F). Cells were prepared following the standard Stellaris RNA FISH protocol for adherent cells (Biosearch Technologies). Cells were fixed using 3.7% formaldehyde in PBS at room temperature for 10 min. Cells were then permeabilized using 70% ethanol for at least 1 h at 4°C. Cells were washed with Wash buffer A and incubated at room temperature for 5 min.

Wash buffer A was aspirated and cells were treated with approximately 30 µl of 125 nM FISH probe diluted in hybridization buffer. The plate was sealed with parafilm and stored in dark humid chamber overnight at 37°C. Cells were then washed with wash buffer A for 30 min at 37°C, then washed with wash buffer B for 5 min and mounted with mounting media containing DAPI (VECTASHIELD #H-2000-10). smFISH imaging was performed using a Zeiss LSM 980 with Airyscan 2 Laser Scanning Confocal imaging system with Zeiss AxioObserver motorized inverted microscope stand, DIC optics, and a motorized XYZ stage. Images were analyzed using IMARIS.

### Western Blots

Samples used for western blots were lysed by boiling in NuPAGE LDS Sample Buffer (Thermo # NP0007) diluted to 1x concentration and supplemented with 1 mM DTT. Samples were run on NuPAGE 4 to 12% Bis-Tris 1.0 mm miniprotein gels (Thermo #NP0321BOX), then transferred to 0.2µM PVDF transfer membranes (Thermo #22860). All protein gels and transfers were run using a mini gel tank (Thermo #A25977) with protein gels run at 200V for 80 min and transfers run at 30V for 60 min. Transferred membranes were blocked in PBS with 1% Triton and 5% milk for at least 1 h at room temperature before they were transferred to primary antibody diluted in the same buffer at 4°C. Primary antibody was applied overnight followed by at least three 5 min washes with the same blocking buffer, secondary antibody diluted in the same buffer for 30 min at room temperature, and three 5 min washes in the same buffer. Westerns were imaged using the Odyssey CLx imaging system by LI-COR. All western blot quantifications were done using Fiji/ImageJ software.

### Polysome profiling

HCT116 cells were cultured in a 15 cm dish, washed twice with cold PBS, scraped into lysis buffer containing 10 mM Tris HCl pH 7.5, 5 mM MgCl_2_, 100 mM KCl, 1% triton, 2 mM DTT, 1x cOmplete mini tablet (EDTA free), and 0.3 U/µl RNasin Plus RNase inhibitor. Cells were lysed by passing through 26G ½ inch needle at least seven times. Lysates were depleted of nuclei by centrifugation at 1300*g* for 10 min at 4°C, and snap frozen using liquid nitrogen and stored at –80°C until use. Samples were loaded onto a 10–50% sucrose gradient and centrifuged at 36,000 rpm for 2 h at 4°C. Polysome profiling was then performed using BioComp Gradient Fractionator. Twelve fractions were collected across approximately 90% of the volume of the sucrose gradient with approximately 800 µl collected per fraction. Each fraction was spiked with 0.1 fmol of an invitro transcribed RNA standard and mixed thoroughly. Samples were then snap frozen using liquid nitrogen and stored at –80°C until later use. RNA extraction and RT-qPCR were then performed on each fraction as described above in the section on tethered reporter TE experiments. Total ribosome loads reported in Fig 6K,N was calculated by multiplying the proportion of reporter RNA from the time course by the mean estimated number of ribosomes found on transcripts in each polysome fraction, and then summing those values across all fractions.

### Ribosome profiling and matched RNA-seq

HCT116 cells were cultured in a 15 cm dish, washed twice with cold PBS, scraped into lysis buffer containing 10 mM Tris HCl pH 7.5, 5 mM MgCl_2_, 100 mM KCl, 1% triton, 2 mM DTT, 1x cOmplete mini tablet (EDTA free), 0.3 U/µl RNasin Plus RNase inhibitor, and 1:60 dilution of ribo-spike. Cells were lysed by passing through 26G ½ inch needle at least 7 times. Lysates were depleted of nuclei by centrifugation at 1300*g* for 10 min at 4°C, divided into aliquots for ribo-seq and matched RNA libraries, and snap frozen using liquid nitrogen. Ribosome profiling and matched RNA seq libraries were then prepared according to previously established protocol,^85^ a detailed version of which is available at http://bartellab.wi.mit.edu/protocols.html. Sequencing was performed on an Illumina HiSeq 2500 using 50 cycles. Only reads mapping to ORFs of GRCh38-annotated genes, fluorescent protein-coding genes, the OsTIR1 gene, or the *fLuc* gene were used. Reads mapping to the first 50 nt of each ORF were excluded. Reads mapping to heme-related genes from the rabbit genome (Broad/oryCun2) were also excluded. TE changes were calculated and count depth normalization was performed for using DESeq2.^86^

### Isolation of RNA from HCT116 cells and SG cores for RNA sequencing

Isolation and sequencing of SG cores was adapted from an established protocol.^18,58^ HCT116 cells were cultured in a 24.5 cm square dish, washed with PBS, scraped into PBS, centrifuged at 1500*g* for 3 min, aspirated to remove liquid, and snap frozen in liquid nitrogen. Frozen cell pellets were thawed in lysis buffer (50 mM Tris HCl pH 7.4, 2 mM MgOAc, 10 mM MgCl_2_, 100 mM KOAc, 0.1% NP40, 0.5 mM DTT, 1x cOmplete mini tablet (EDTA free), and 1 U/µl RNasin Plus RNase inhibitor). Cells were lysed by passing through 26G ½ inch needle at least seven times.

Lysates were depleted of their nuclei by centrifugation at 1000*g* for 5 min at 4°C. 5% of this nuclear-depleted sample was aliquoted and snap frozen in liquid nitrogen and stored at –80°C as the cytoplasmic fraction. The remaining 95% was used to isolate the SG fraction. Sample was centrifuged at 18,000*g* for 20 min at 4°C. Pellet was resuspended in 1 ml lysis buffer and allowed to turn end-over-end for at least 10 min at 4°C. Once resuspended, sample was centrifuged again at 18,000*g* for 20 min at 4°C. The pellet was resuspended in 300 µl lysis buffer and centrifuged at 850*g* for 2 min at 4°C. The supernatant was taken and incubated with 6 µg of G3BP2 antibody (Bethyl Laboratories #A302-040A) overnight at 4°C, rotating end-over-end.

Sample was centrifuged at 18,000*g* for 20 min at 4°C to remove unbound antibody, and the pellet was resuspended in 500 µl lysis buffer. 625 µl of M-280 sheep, anti-rabbit IgG dynabeads (Thermo # <u>11204D</u>) were then added to the sample and incubated for at least 3 h at 4°C, rotating end-over-end. Beads were washed with buffer containing 20 mM Tris HCl pH 8.0 and 50 mM NaCl and eluted using a high-salt buffer (10mM Tris HCl pH 7.5, 1M NaCl, 5% 1,6 hexanediol, 5mM MgCl_2_, 100mM KCl, 1% triton, 2mM DTT, 1x cOmplete mini tablet (EDTA free), and 0.3 U/µl RNasin Plus RNase inhibitor), turning end-over-end for at least 4 h. Eluate was snap frozen using liquid nitrogen and stored at –80°C as the SG fraction. RNA-sequencing libraries were then prepared from the cytoplasmic and SG fractions in parallel, following the protocol used for the RNA-sequencing libraries that matched the ribosome profiling. SG enrichment values were calculated using DESeq2.^86^

## Data availability

Raw sequencing data for all ribo-seq and SG purification experiments of this study have been deposited in the Gene Expression Omnibus (GEO) under accession code GSE256237. Previously published sequencing data and re-analyzed using R and are available under accession codes GSE99304, GSE131650, GSE254636, GSE254637, GSE254638, GSE90869, GSE103667, and GSE223295.^18,42,55,59,60^ Source data are provided with this study. All other data supporting the findings of this study are available from the corresponding author on reasonable request.

## Statistical analysis and comparison with published datasets

Plots and statistical analyses were performed using R. No statistical methods were used to pre-determine sample sizes but our sample sizes are similar to those reported in previous publications.^18,55,59^ Statistical tests and parameters are shown in figure legends. Data distribution was assumed to be normal but this was not formally tested. Data collection and analysis were not performed blind to the conditions of the experiments. Additional data were obtained from publicly available datasets associated with published work and re-analyzed using R.^18,42,55,59,60^

## Author Contributions

J. S. conceived the project and designed the experiments. J. S. performed the experiments and analyses. J. S. and D. B. designed the figures and wrote the paper. The paper was read and approved by all authors.

## Acknowledgements

We thank members of the Bartel laboratory for helpful discussions, R. Young at the Whitehead Institute for providing HCT116 cells, the Whitehead Genome Technology Core for high-throughput sequencing, and the Whitehead Flow Cytometry Core for cell sorting. J. S. is a Hanna Gray Fellow of the Howard Hughes Medical Institute; D. B. is an investigator of the Howard Hughes Medical Institute.

---

**Extended Data Figure 1.**
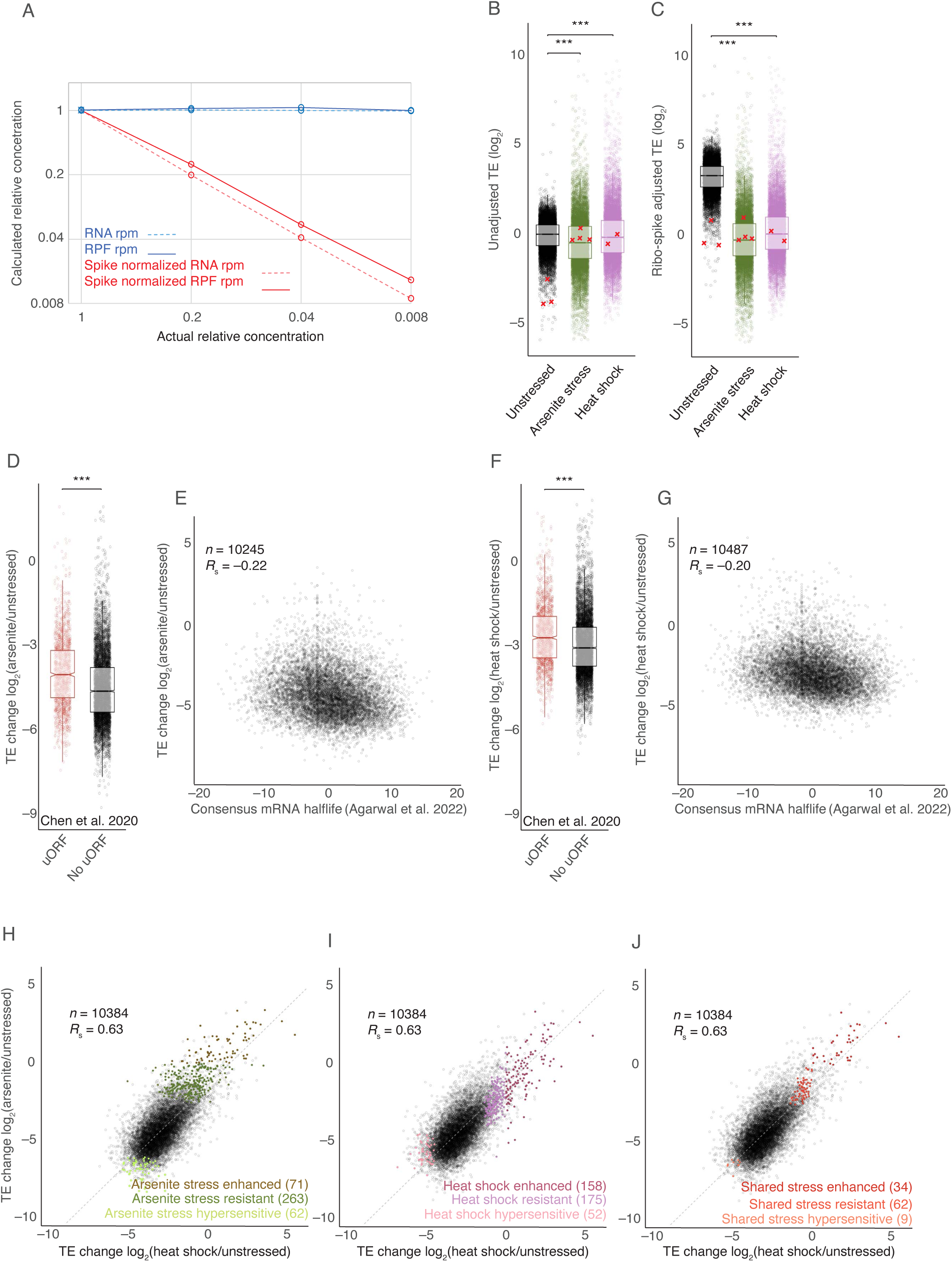
Heat shock and oxidative stress illicit similar but distinct ISR translation programs. **(A)** Quantification of ribo-spike. To confirm that ribo-spikes functioned as intended, cytoplasmic extract from HCT116 cells was serially diluted to achieve the indicated relative concentrations, and then a constant amount of ribo-spike was added to each sample before ribosome profiling and RNA-seq analyses. Plotted are the relative concentrations of total mRNA (dashed line) and RPFs (solid line) as calculated from the 1000 most highly expressed genes, using no-spike controls (blue) and ribo-spike normalized samples (red) as a function of the actual relative concentrations. Spike-normalized RPF and mRNA concentrations closely matched the known dilutions. **(B)** Relative TE measurements before adjusting for ribo-spike. Plotted with circles are the average log_2_ TE values (line, median; notch, 95% confidence interval; box, quartiles; whiskers, 1.5 x IQR) calculated for mRNAs from either untreated cells (black, three replicates), cells treated with 500 µM sodium arsenite for 1 h (green, four replicates), or cells treated with heat shock at 45°C for 25 min (pink, 2 replicates). Red Xs denote the ribo-spike TE values (one for each replicate). Statistical significance was calculated based on ribo-spike values alone (***, *p* ≤ 0.001; Welch’s two-sample t-test). **(C)** TE measurements of panel B after adjusting for ribo-spike; otherwise, as in (B). Note that statistical significance is determined in panel B, using ribo-spike values prior to adjustment (***, *p* ≤ 0.001). **(D)** Preferential translation of mRNAs containing uORFs during arsenite stress. The plots indicate log_2_ FC in TE caused by arsenite for mRNAs containing at least one uORF and those containing no uORFs, according to annotations from Chen et al. (2020)^42^ (line, median; notch, 95% confidence interval; box, quartiles; whiskers, 1.5 x IQR); ***, *p* ≤ 0.001; Welch’s two-sample t-test). **(E)** Relationship between translation during arsenite stress and mRNA half-lives. The plots indicate the log_2_ FC in TE due to arsenite as a function of the mean-normalized half-life (PC1) of mRNAs as calculated from consensus of multiple human cell data sets by Agarwal et al. (2022).^45^ *n* indicates number of unique mRNAs. **(F)** Preferential translation of mRNAs containing uORFs during heat shock. Plots indicate log_2_ FC in TE caused by treatment with 45°C heat shock; otherwise, as in (D). **(G)** Relationship between translation during heat shock and mRNA half-lives. The plots indicate the log_2_ FC in TE due to heat shock as a function of the mean-normalized half-life of mRNAs; otherwise, as in (E). **(H – J)** Comparison of arsenite stress and heat shock ISR translation programs. Plotted for each mRNA are changes in TE caused by arsenite as a function of changes in TE caused by heat shock. Points for mRNAs classified as either stress-enhanced, stress-resistant, or stress-hypersensitive mRNAs in either arsenite stress (D), heat shock (E), or both (F) are colored. *n* indicates number of unique mRNAs; *R*_s_ indicates Spearman coefficient.

**Extended Data Figure 2.**
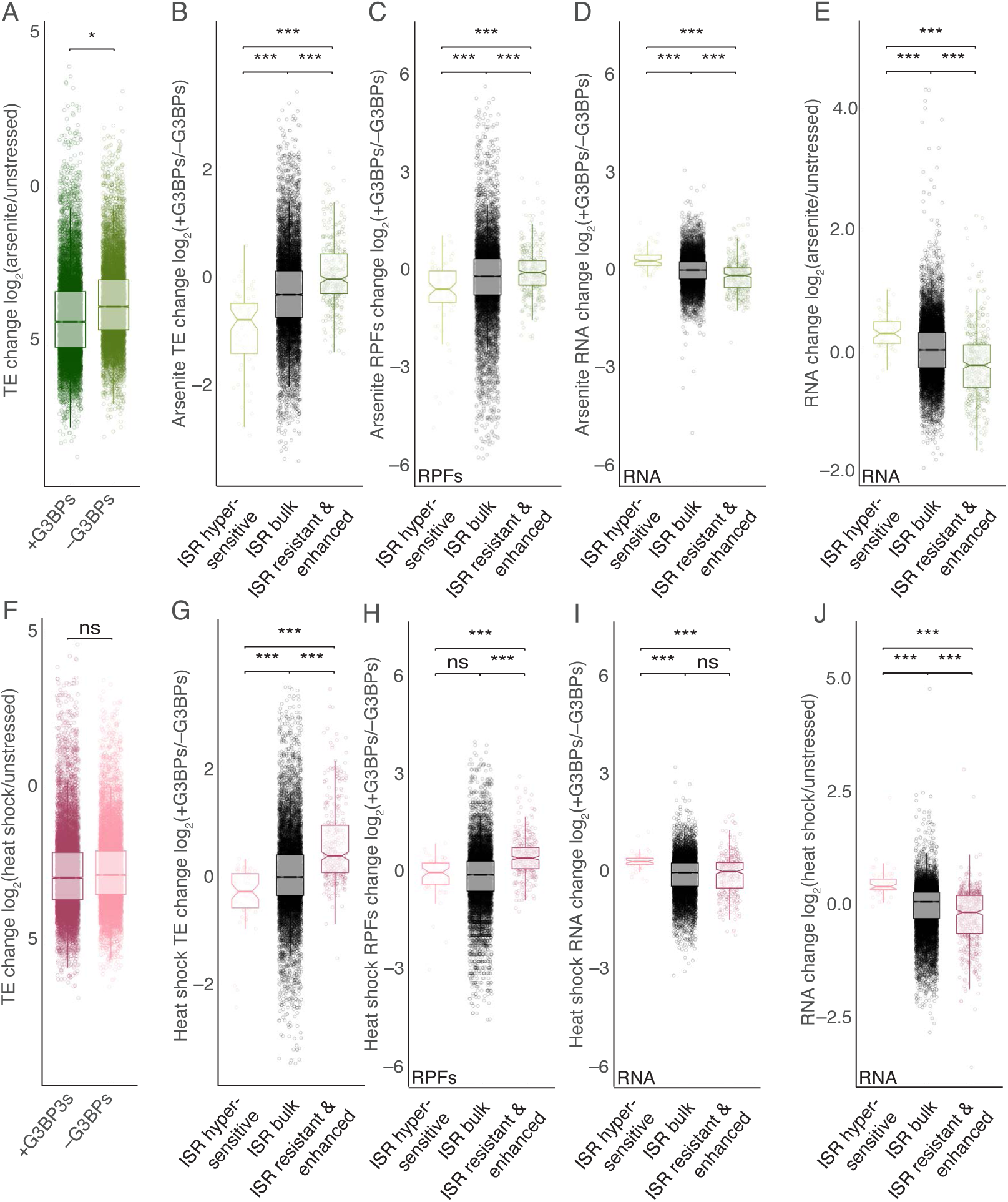
G3BPs reinforce the ISR translation program across multiple cell types and stresses. **(A)** The ISR translation response to arsenite stress, either with or without G3BPs. The plots indicate the log_2_ FC in TE caused by arsenite stress in either untreated HCT116 cells or cells depleted of their G3BPs by addition of 500 µM IAA (line, median; notch, 95% confidence interval; box, quartiles; whiskers, 1.5 x IQR). Significance was determined using ribo-spike values, as in Figure S1B (* *p* ≤ 0.05, ***, *p* ≤ 0.001; Welch’s two-sample t-test). **(B)** Comparison of G3BP-dependent translation and ISR translation in HCT116 cells treated with arsenite stress. The plots indicate the log_2_ FC in TE due to presence of G3BPs in stressed cells in each of three ISR translation categories: stress-enhanced/resistant, bulk behavior, and stress-hypersensitive (line, median; notch, 95% confidence interval; box, quartiles; whiskers, 1.5 x IQR; ***, *p* ≤ 0.001; Welch’s two-sample t-test). **(C)** Comparison of G3BP-dependent RPF changes in HCT116 cells treated with arsenite stress. The plots indicate the fold change in RPFs due to presence of G3BPs in stressed cells; otherwise, as in (B). **(D)** Comparison of G3BP-dependent RNA changes in HCT116 cells undergoing arsenite stress. The plots indicate the fold change in RNA due to presence of G3BPs in stressed cells; otherwise, as in (B). **(E)** Comparison of stress-dependent RNA changes in HCT116 cells undergoing arsenite stress. The plots indicate the fold change in RNA due to treatment with 500 µM sodium arsenite for 1 h; otherwise, as in (B). **(F)** The ISR translation response to heat shock, either with or without G3BPs. ns, not significant; otherwise, as in (A). **(G)** Comparison of G3BP-dependent translation and ISR translation in HCT116 cells undergoing heat-shock stress; otherwise, as in (B). **(H)** Comparison of G3BP-dependent RPF changes in HCT116 cells undergoing heat-shock stress. The plots indicate the fold change in RPFs due to presence of G3BPs in stressed cells; otherwise, as in (B). **(I)** Comparison of G3BP-dependent RNA changes in HCT116 cells undergoing heat-shock stress. The plots indicate the fold change in RNA due to presence of G3BPs in stressed cells; otherwise, as in (B). **(J)** Comparison of stress-dependent RNA changes in HCT116 cells undergoing heat-shock stress. The plots indicate the fold change in RNA due to treatment with heat-shock stress; otherwise, as in (B).

**Extended Data Figure 3.**
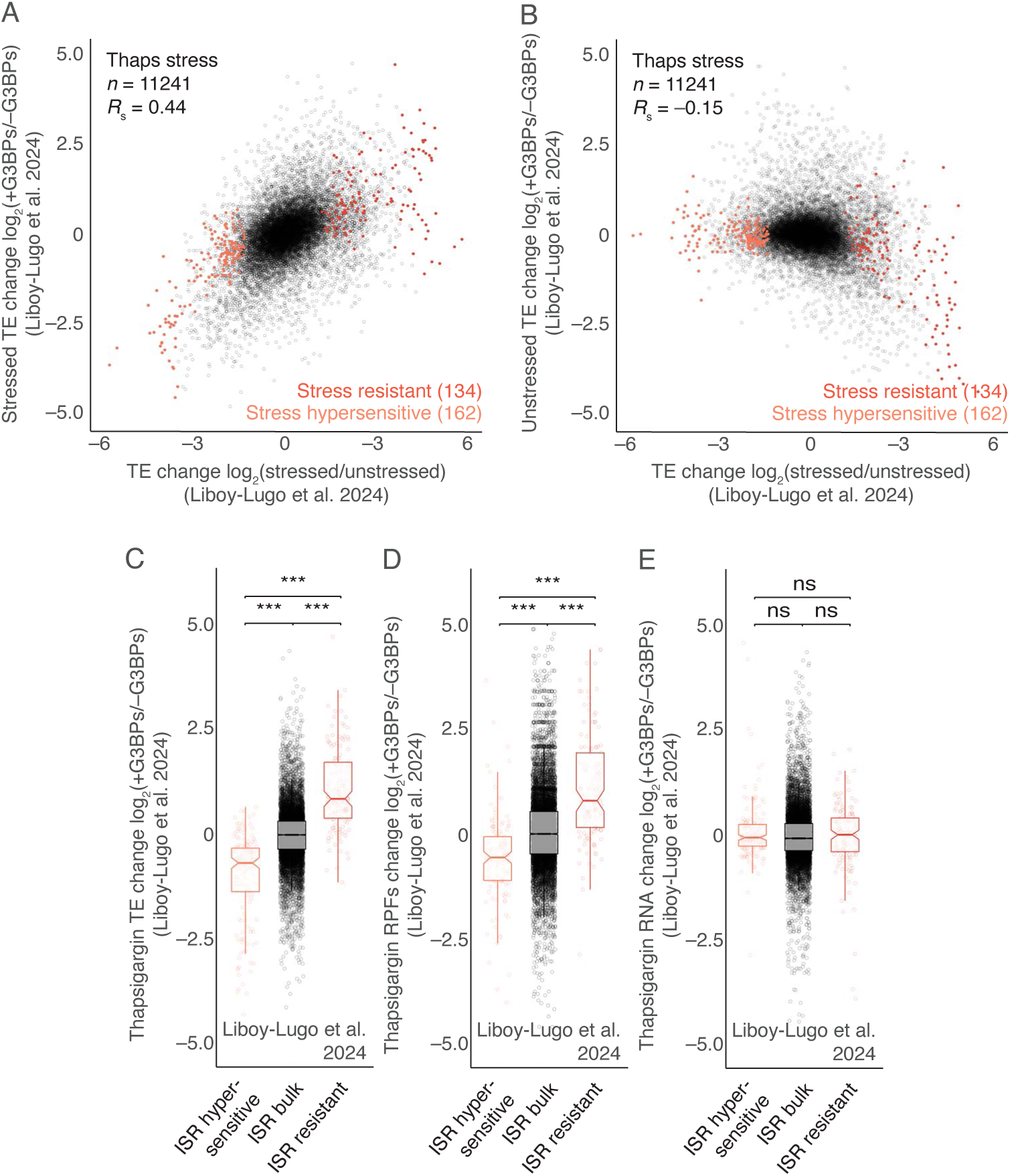
G3BPs reinforce the ISR translation program across multiple cell types and stresses. **(A)** Relationship between G3BP-dependent translation during thapsigargin stress and the ISR translation program as measured by Liboy-Lugo et al. (2024);^55^ otherwise as in Fig 2H. **(B)** Relationship between G3BP-dependent translation in the absence of stress and the ISR translation program as measured by Liboy-Lugo et al. (2024);^55^ otherwise as in Fig 2I. **(C)** Comparison of G3BP-dependent translation and ISR translation in U2OS cells undergoing thapsigargin stress as measured by Liboy-Lugo et al. (2024);^55^ otherwise as in (Fig S2B). **(D)** Comparison of G3BP-dependent RPF changes in U2OS cells undergoing thapsigargin stress as measured by Liboy-Lugo et al. (2024);^55^ otherwise as in (Fig S2C). **(E)** Comparison of G3BP-dependent RNA changes in U2OS cells undergoing thapsigargin stress as measured by Liboy-Lugo et al. (2024);^55^ otherwise as in (Fig S2D).

**Extended Data Figure 4.**
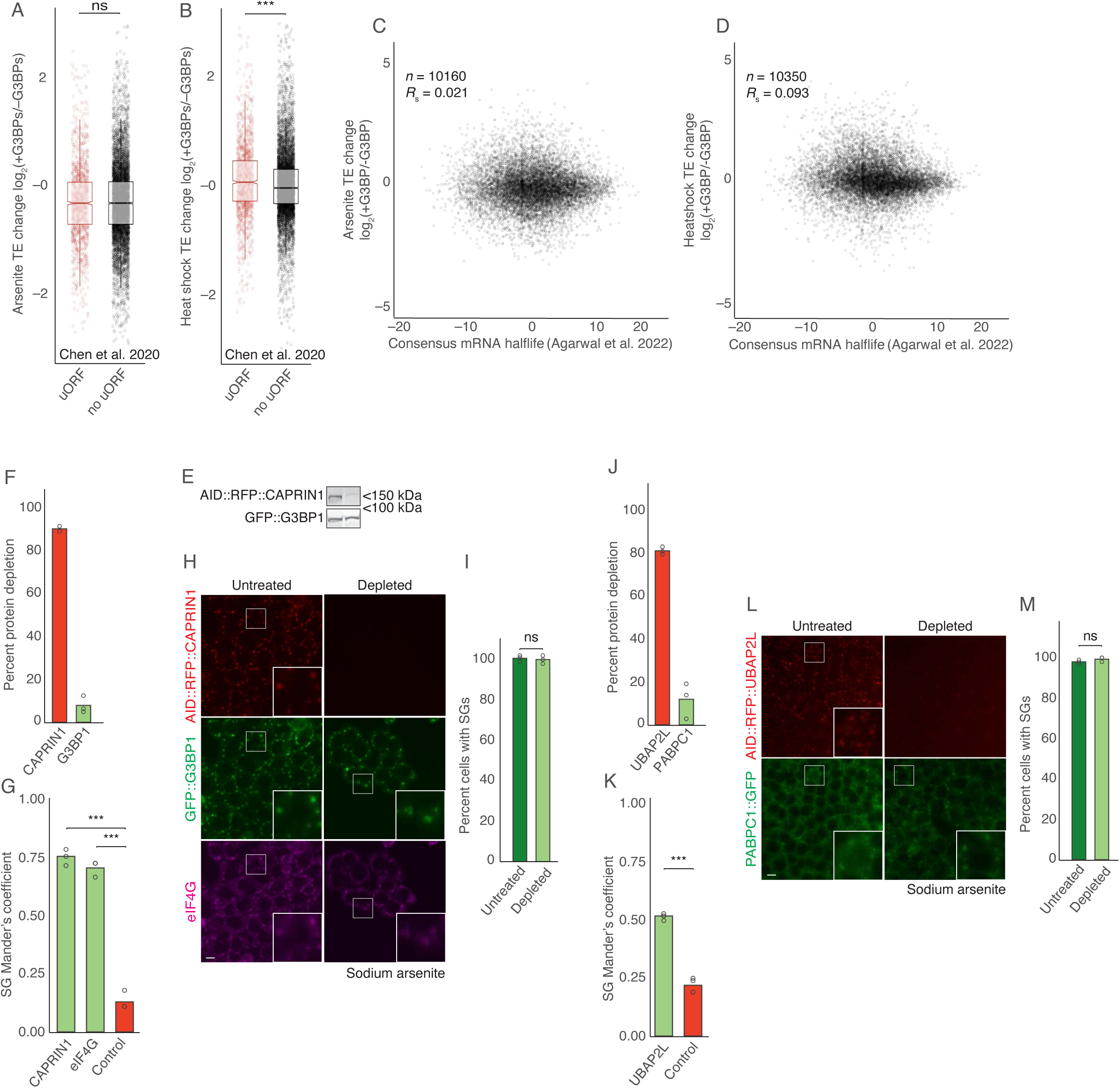
Neither CAPRIN1 nor UBAP2L are required for SG formation in HCT116 cells. **(A)** G3BP-dependent translation of mRNAs containing uORFs during arsenite stress. The plots indicate the log_2_ FC in TE due to presence of G3BPs in arsenite-stressed cells for mRNAs containing at least one uORF and those containing no uORFs, according to annotations from Chen et al. (2020)^42^ (line, median; notch, 95% confidence interval; box, quartiles; whiskers, 1.5 x IQR); ***, *p* ≤ 0.001; Welch’s two-sample t-test). **(B)** G3BP-dependent translation of mRNAs containing uORFs during heat stress. The plots indicate the log_2_ FC in TE due to presence of G3BPs in heat-stressed cells for mRNAs containing at least one uORF and those containing no uORFs. We observed that mRNAs containing one or more functional uORF were preferentially translated in the presence of heat stress. However, as this trend did not hold in arsenite-stressed cells, suggesting that uORFs might play only a minor or a stress-specific role in G3BP-dependent translation; otherwise as in (A). **(C)** Relationship between G3BP-dependent translation during arsenite stress and mRNA half-lives. We observed no correlation between the half-life of an mRNA and its G3BP-dependent translation, indicating that preferential translation of newly synthesized mRNAs might be stress-granule independent. The plots indicate the log^2^ FC in TE due to presence of G3BPs in arsenite-stressed cells as a function of the mean-normalized half-life (PC1) of mRNAs as calculated from consensus of multiple human cell data sets by Agarwal et al. (2022).^45^ *n* indicates number of unique mRNAs. **(D)** Relationship between G3BP-dependent translation during heat stress and mRNA half-lives. The plots indicate the log_2_ FC in TE due to presence of G3BPs in heat-stressed cells as a function of the mean-normalized half-life of mRNAs; otherwise, as in (C). **(E)** Depletion of endogenous CAPRIN1 fused to AID. Immunoblots probed for the indicated proteins show specific depletion of AID fusion protein after treatment with 500 µM IAA for 3 h. **(F)** Quantification of protein depletion in (E), normalizing to levels of GAPDH. Points show values for three biological replicates. **(G)** Colocalization of CAPRIN1 in SGs. Quantification of colocalization was performed using Mander’s coefficient between G3BP1 and indicated SG marker, comparing to a control image from an irrelevant field of cells (***, *p* ≤ 0.001; Welch’s two-sample t-test)**. (H)** SG formation despite CAPRIN1 depletion. HCT116 cells were either treated with 500 µM IAA to deplete CAPRIN1 (right) or not treated (left) prior to arsenite stress (500 µM sodium arsenite for 1 h). Images show either CAPRIN1 in red, SG marker protein G3BP1 in green, or SG marker protein eIF4G in magenta (scale bars, 10 µm). **(I)** Quantification of SG formation in (P). Points show values for three biological replicates (ns, not significant). **(J)** Depletion of endogenous UBAP2L fused to AID. Quantification was based on AID::RFP::UBAP2L and PABPC1::GFP fluorescence in homozygously tagged HCT116 cells (images not shown). Points show values for three biological replicates. **(K)** Colocalization of UBAP2L in SGs. Quantification of colocalization was performed using Mander’s coefficient between PABPC1 and UBAP2L, comparing to a control image from an irrelevant field of cells (***, *p* ≤ 0.001; Welch’s two-sample t-test)**. (L)** SG formation despite UBAP2L depletion. HCT116 cells were either treated with 500 µM IAA to deplete UBAP2L (right) or not treated (left) prior to arsenite stress (500 µM sodium arsenite for 1 h). Images show either UBAP2L in red or SG marker protein PABPC1 in green; otherwise, as in (P). **(M)** Quantification of SG formation in (L); otherwise, as in (I).

**Extended Data Figure 5.**
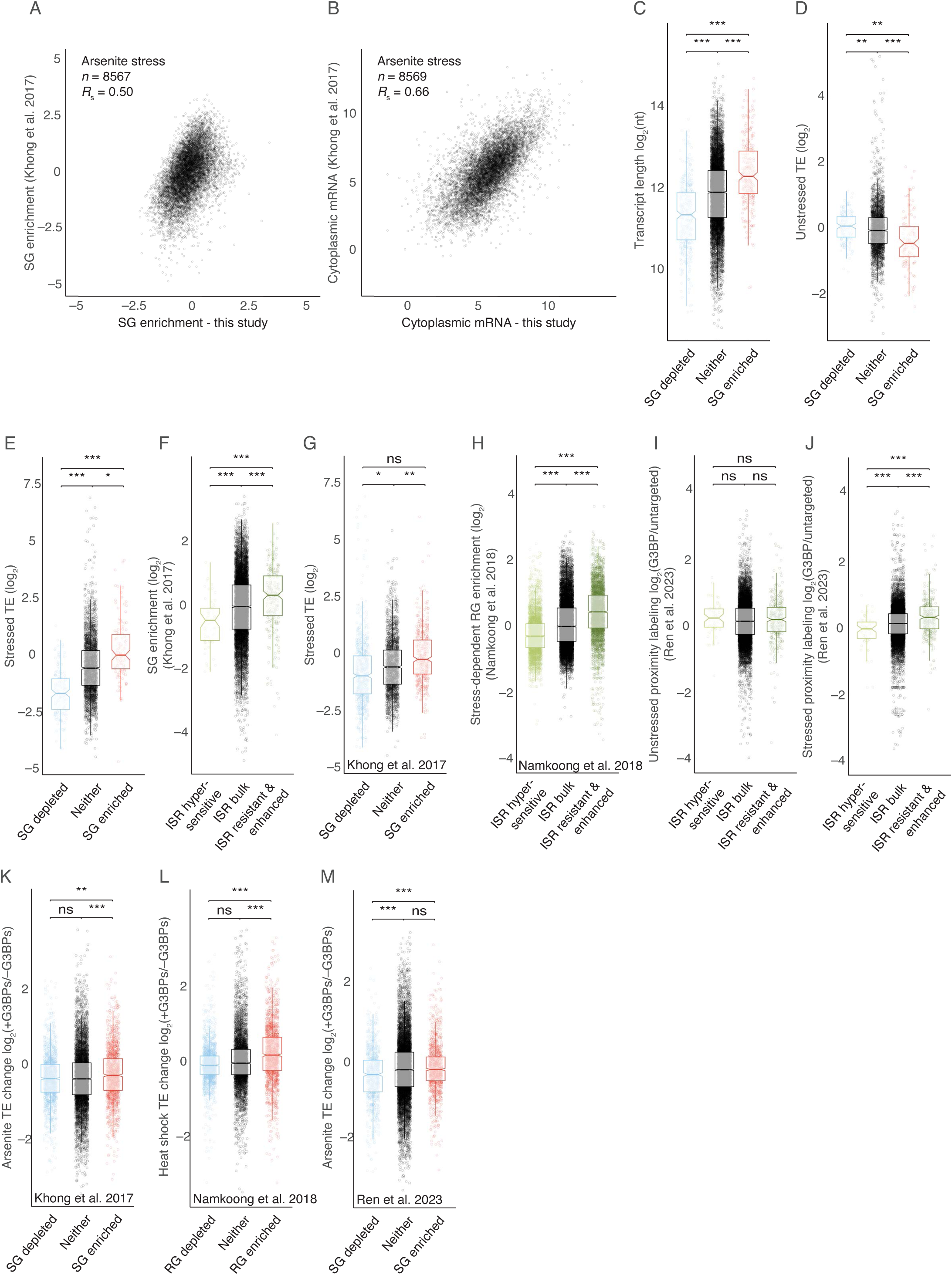
Conclusions regarding SG-enrichment are consistent across multiple cell types and stresses. **(A)** Comparison of SG enrichments. Shown are SG enrichments calculated in Khong et al. (2017)^18^ plotted as a function of SG enrichments calculated in this study. *n* indicates total number of unique mRNAs. **(B)** Comparison of mRNA levels for the cells used in (A). Shown are relative mRNA abundances in U2OS cells, as measured by Khong et al. (2017)^18^, plotted as a function of relative abundance in HCT116 cells, as measured in this study. Otherwise as in (A). Our HCT116 cell SG enrichments resembled those previously reported in U2OS cells, especially when considering the differences in overall mRNA expression in the two cell types. **(C – E)** SG enrichment determinants. Shown is the relationship between either transcript length (C), unstressed TE (D), or stressed TE (E) and SG enrichment. The relationship between ISR translation change and SG enrichment was not simply a consequence of SGs being enriched for transcripts that were poorly translated prior to stress and therefore experienced a smaller decline, as SGs were also enriched for mRNAs that were translated relatively well during stress, as shown in E. otherwise as in Fig 3C (* *p* ≤ 0.05, ** *p* ≤ 0.01, *** *p* ≤ 0.001). **(F)** Comparison of SG enrichment in arsenite-stressed U2OS cells as measured by Khong et al. (2017)^18^ and ISR translation in arsenite stressed HCT116 cells; otherwise, as in Fig 3B. **(G)** Comparison of arsenite-stressed TE in HCT116 cells and SG enrichment in arsenite-stressed U2OS cells as measured by Khong et al. (2017)^18^; otherwise, as in (E). **(H)** Comparison of Stress-Dependent RNP Granule (RG) enrichment and ISR translation in ER-stressed NIH 3T3 cells as measured by Namkoong et al. (2018);^59^ otherwise, as in (F). **(I)** Comparison of G3BP-depdenent proximity labeling in unstressed HEK293T cells as measured by Ren et al. (2023)^60^ and ISR translation in arsenite stressed HCT116 cells. ns, not significant; otherwise, as in (F). **(J)** Comparison of G3BP-depdenent proximity labeling in arsenite-stressed HEK293T cells as measured by Ren et al. (2023)^60^ and ISR translation in arsenite-stressed HCT116 cells; otherwise, as in (I). **(K)** Comparison of G3BP-dependent translation in HCT116 cells and SG enrichment in U2OS cells as measured by Khong et al. (2017)^18^. All stresses were 500 µM sodium arsenite for 1 h; otherwise, as in (E). **(L)** Comparison of G3BP-dependent translation in HCT116 cells and RG enrichment in NIH 3T3 cells as measured by Namkoong et al. (2018).^59^ All stresses were heat shock; otherwise, as in (I). **(M)** Comparison of G3BP-dependent translation in HCT116 cells and G3BP-depdenent proximity labeling in stressed HEK293T cells as measured by Ren et al. (2023)^60^. All stresses were 500 µM sodium arsenite for 1 h; otherwise, as in (K).

**Extended Data Figure 6.**
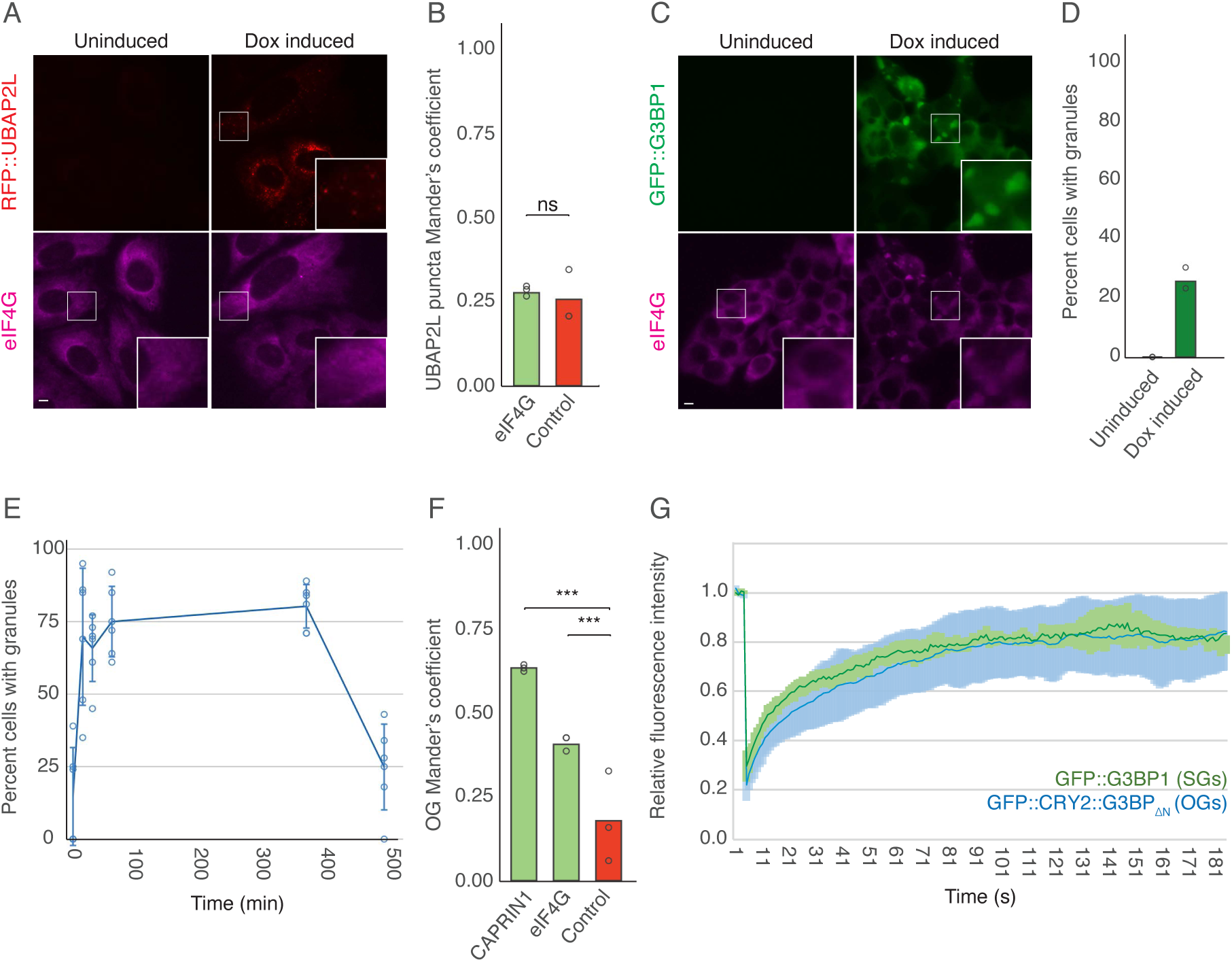
OGs are SG-like condensates formed in the absence of stress. **(A)** No induction of ectopic SGs caused by UBAP2L overexpression. Unstressed U2OS cells were either treated with 1 µM doxycycline (right) or not treated (left) to induce overexpression of UBAP2L. Images show either UBAP2L in red or SG marker protein eIF4G in magenta (scale bars, 10 µm). White boxes highlight insets that are expanded on the right. **(B)** Colocalization of UBAP2L puncta with SG marker. Quantification of colocalization was performed using Mander’s coefficient between UBAP2L and eIF4G as an SG marker, comparing to a control image from an irrelevant field of cells (***, *p* ≤ 0.001; Welch’s two-sample t-test)**. (C)** Induction of ectopic SGs caused by G3BP overexpression. Unstressed HCT116 cells were either treated with 1 µM doxycycline (right) or not treated (left) to induce overexpression of G3BP1. Images show either G3BP1 in green or SG marker protein eIF4G in magenta; otherwise, as in (A). **(D)** Quantification of ectopic granule formation in (C). Points indicate values for three biological replicates. **(E)** Kinetics of OG formation and dissolution. Plotted over time is the percent of cells with visible OGs. Cells were exposed to blue light starting at 0 min and removed from blue light at 360 min. Error bars report standard deviation for six biological replicates shown as points. **(F)** Colocalization of SG markers in OGs. Quantification of colocalization was performed using Mander’s coefficient between GFP::CRY2::G3BP_ΔN_ and indicated SG marker, comparing to a control image from an irrelevant field of cells (***, *p* ≤ 0.001; Welch’s two-sample t-test)**. (G)** FRAP of OGs. Plotted over time is the fluorescence intensity of GFP-labeled granules after photobleaching. SG (green) and OG (blue) recoveries are shown over a 180 s time course. Photobleaching occurred at 3 s. Error bars report standard deviation for at least six granules from at least three cells.

**Extended Data Figure 7.**
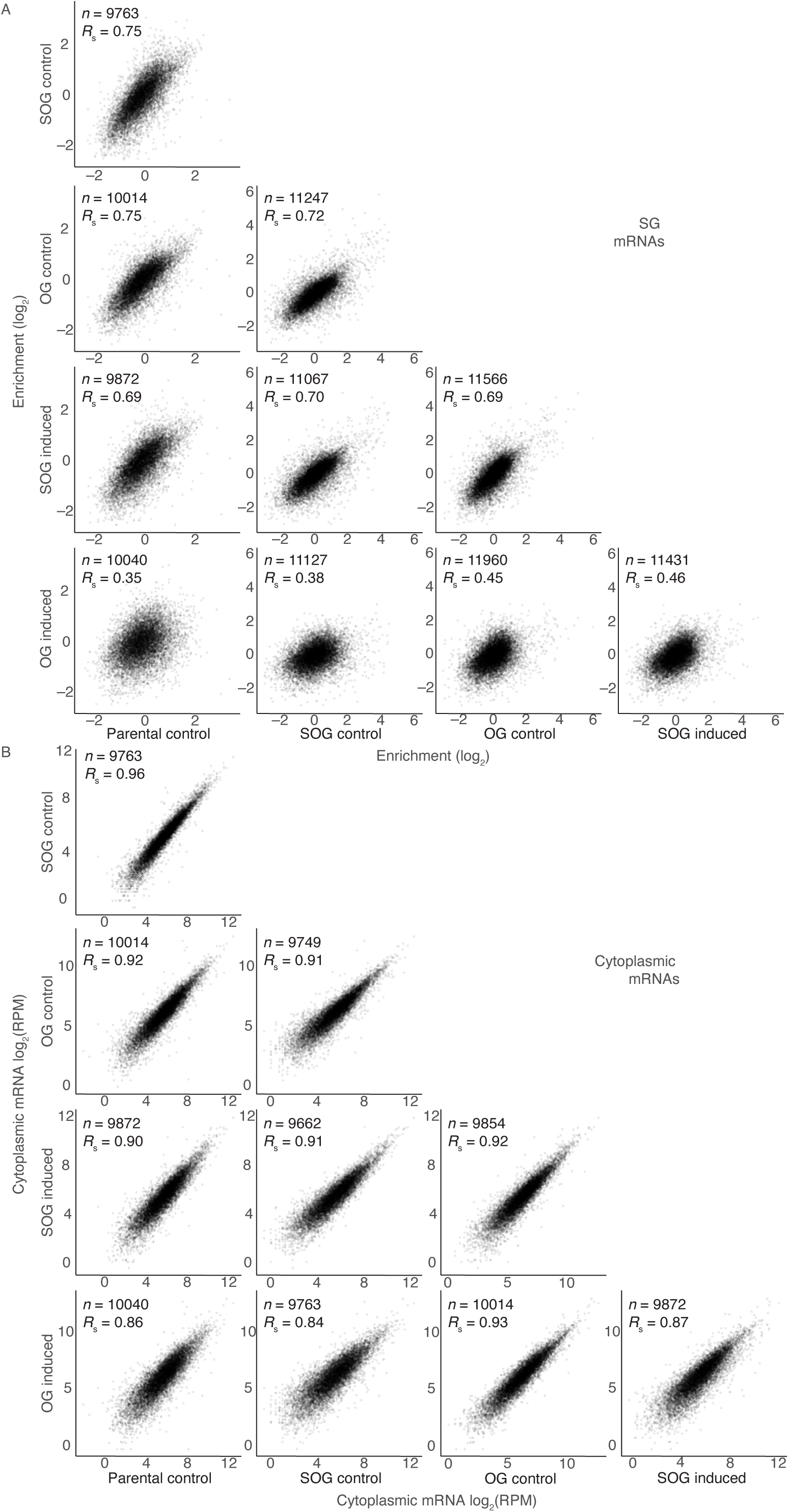
OGs require stress to recruit the SG transcriptome. **(A)** Expanded comparisons of granule enrichment, showing the distributions used to calculate the *R*_S_ values of Fig 5E. *n* indicates total number of unique mRNAs. **(B)** Expanded comparisons of cell line transcriptomes, showing the distributions used to calculate the *R*_S_ values of Fig 5F.

**Extended Data Figure 8.**
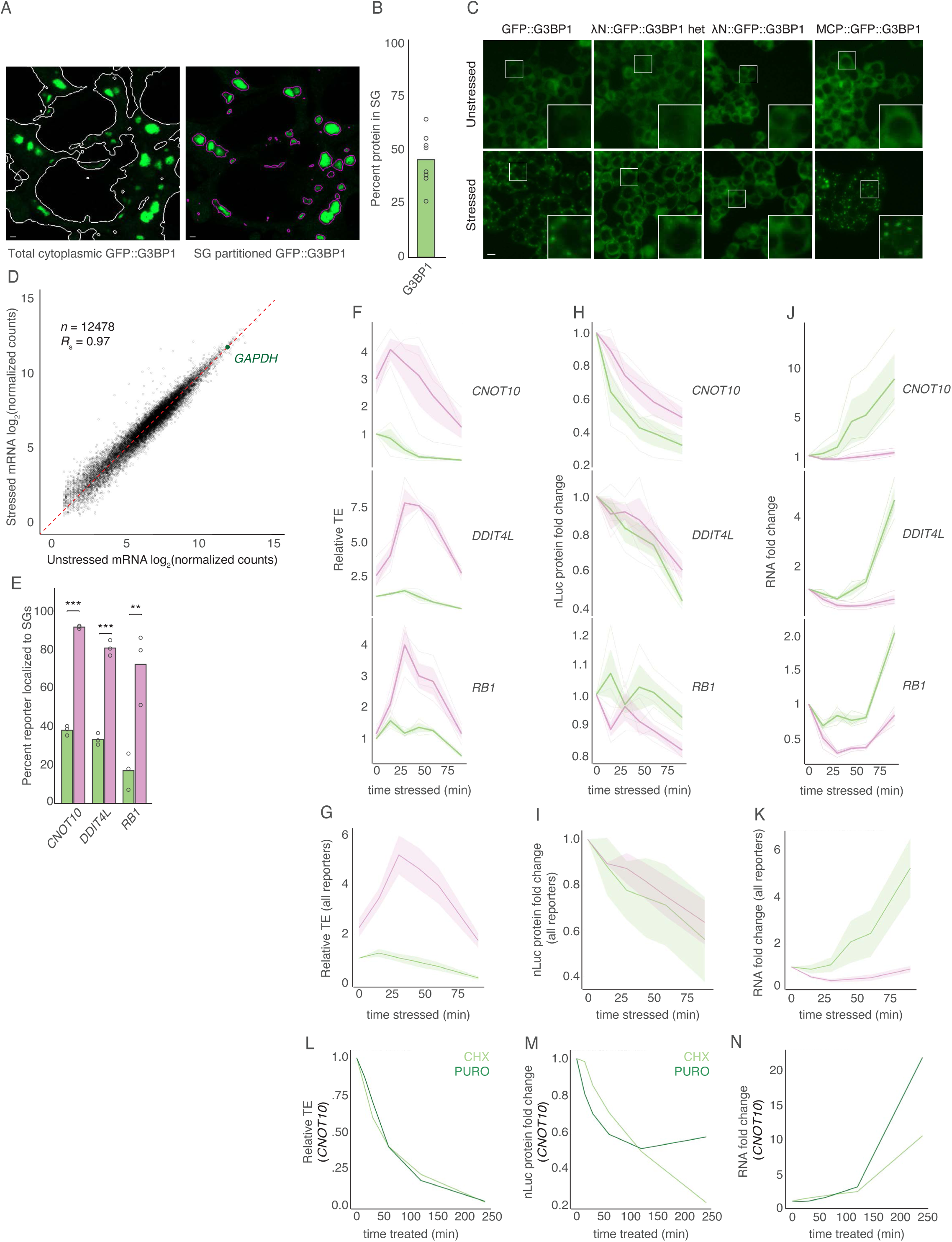
G3BP1-tethered preferential translation occurs across multiple reporters and correlates with translation-dependent mRNA destabilization. (**A**) G3BP1 partitioning to SGs. HCT116 cells expressing G3BP1 endogenously tagged with GFP were stressed with 500 µM sodium arsenite for 90 min. Images show localization of G3BP in green. White outlines show cell boundaries as determined by Imaris software based on cytosolic fluorescence. Purple outlines show SG boundaries as determined by Imaris software (scale bars, 1 µm). (**B**) Quantification of G3BP1 partitioning to SGs. Points show the amount of fluorescence from SGs as a percent of total fluorescence, for eight independent replicates each containing at least three cells. (**C**) Effects of tagging G3BP with λN and MCP. HCT116 cells expressing endogenous G3BP tagged with the indicated GFP fusion protein were either unstressed (top) or treated with 500 µM sodium arsenite for 1 h (bottom) to assess SG formation (scale bar, 10 µm). White boxes highlight insets that are expanded on the right. (**D**) mRNA changes during stress. Plotted for each mRNA is its abundance in arsenite-stressed HCT116 cells (500 µM sodium arsenite for 1 h) as a function of its abundance in unstressed HCT116 cells. The *GAPDH* mRNA, which was used as a control for RT-qPCR, is highlighted in green. The red dashed line indicates unchanged abundance. *n* indicates total number of unique mRNAs. **(E)** SG localization for each reporter. Percent of cytoplasmic reporter molecules localized to SGs in untethered cells (green) and in tethered cells (pink) is shown for each reporter with 5’ and 3’ UTR sequences labeled below. Points show values for at least three biological replicates (** *p* ≤ 0.01, *** *p* ≤ 0.001, Welch’s two-sample t-test). The average SG tethering efficiency for the three different reporters was 30% for untethered cells and 82% for tethered cells, with standard deviations of 11% and 10%, respectively. **(F)** G3BP1-tethered ISR translation response for all reporters, plotted as in Fig 6D. For comparison, results from Fig 6D are replotted. **(G)** Average ISR translation response across all G3BP1-tethered reporters. Plotted are the average TE values across all three reporters shown in (F). **(H)** G3BP1-tethered ISR protein response for all reporters. Plotted is the fold change in luciferase protein in untethered cells (green) and tethered cells (pink) during a time course of treatment with arsenite stress (500 µM sodium arsenite). Luciferase levels are reported relative to those of unstressed cells from the same biological replicate; otherwise as in (F). **(I)** Average ISR protein response across all G3BP1-tethered reporters. Plotted are the average luciferase values across all three reporters shown in (H). **(J)** G3BP1-tethered ISR RNA response for all reporters. Plotted is the fold change in reporter mRNA in untethered cells (green) and tethered cells (pink) during a time course of treatment with arsenite stress (500 µM sodium arsenite). RNA levels are reported relative to those of unstressed cells from the same biological replicate; otherwise as in (F). **(K)** Average ISR RNA response across all G3BP1-tethered reporters. Plotted are the average mRNA fold changes across all reporters shown in (J). **(L)** Reporter translation response to translational inhibitors. Plotted is the TE of the *CNOT10* reporter in untethered cells treated with 100 µg/ml cycloheximide (light green) or 100 µg/ml puromycin (dark green) in a time course. Time points were taken at 0, 15, 30, 60, 120, and 240 min; otherwise as in (F). **(M)** Reporter protein response to translational inhibitors. Plotted is the fold change in luciferase protein of the *CNOT10* reporter during the time course shown in (L); otherwise as in (G).

**Extended Data Figure 9.**
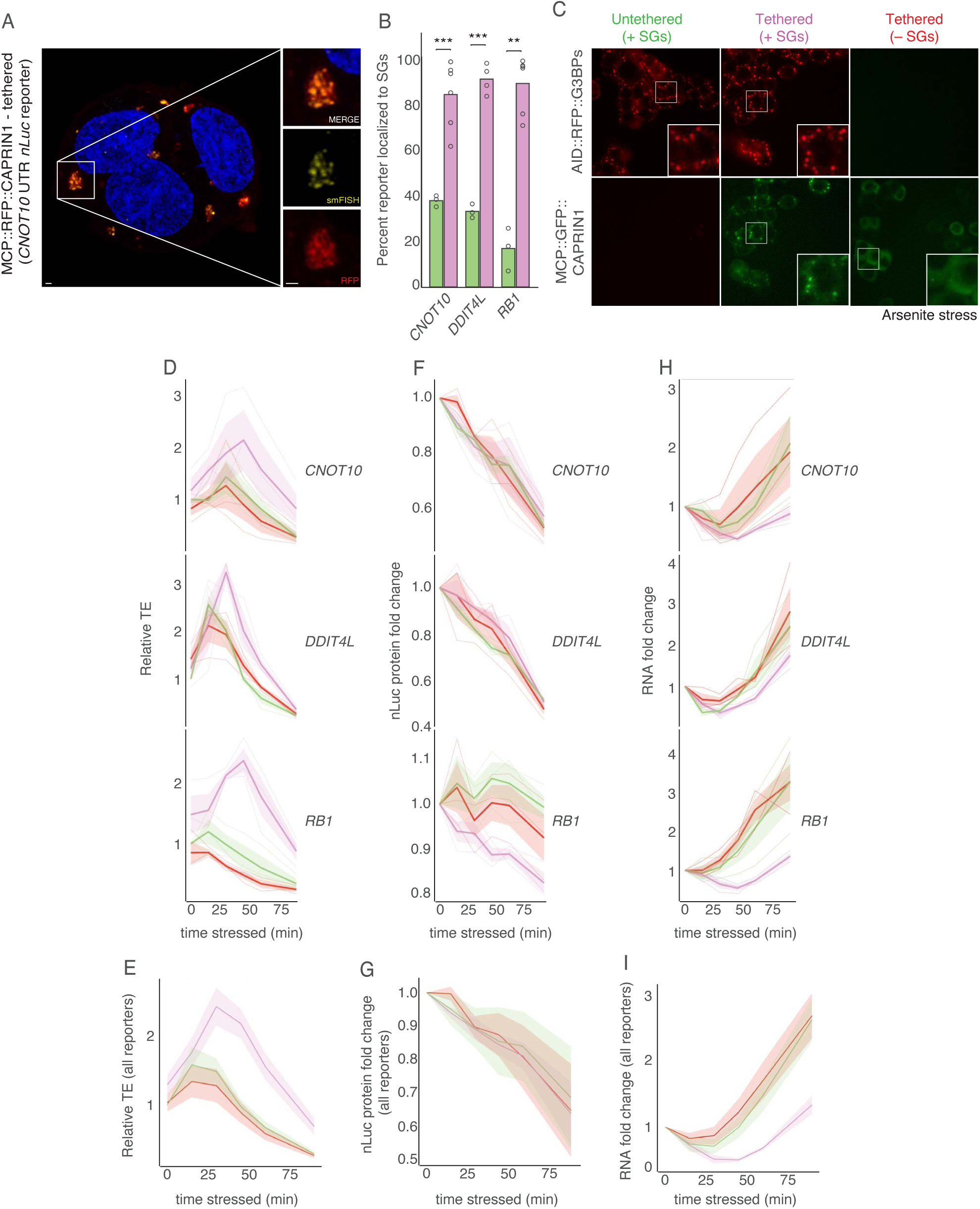
CAPRIN1-tethered preferential translation occurs across multiple reporters and correlates with translation-dependent mRNA destabilization. **(A)** Tethering a luciferase reporter to SGs via CAPRIN1. HCT116 cells expressing a NanoLuc reporter bearing 24x MS2 hairpins and the 5’ and 3’ UTRs from *CNOT10* along with MCP::RFP::CAPRIN1 were stressed with 500 µM sodium arsenite for 90 min. Images show localization of CAPRIN1 in red and smFISH of NanoLuc reporter molecules in yellow (scale bars, 1 µm). White boxes highlight insets that are expanded on the right. **(B)** CAPRIN1 tethering efficiency. Percent of cytoplasmic reporter molecules localized to SGs in untethered cells (replotted from Figure S6E in green) and to SGs in CAPRIN1-tethered cells (pink) are shown for reporters with 5’ and 3’ UTRs of the indicated genes. Points show values for at least three biological replicates (** *p* ≤ 0.01, *** *p* ≤ 0.001, Welch’s two-sample t-test). The average SG tethering efficiency for the four different reporters was 30% for untethered cells and 89% for tethered cells, with standard deviations of 11% and 4%, respectively. **(C)** Assessment of SG formation in cells expressing CAPRIN1 tether while depleted of G3BPs. HCT116 cells expressing endogenously tagged AID::RFP::G3BPs were either untreated (left), treated with 1 µM doxycycline and 200 µM auxinole for 10 h, leading to retention of G3BPs and expression of MCP::GFP::CAPRIN1 (middle), or treated with 1 µM doxycycline and 500 µM IAA for 10 h, leading to turnover of endogenous G3BPs and expression of MCP::GFP::CAPRIN1 (right). Cells were subsequently stressed with 500 µM sodium arsenite for 90 min. White boxes highlight insets that are expanded on the right. Scale bar, 10 µm. These results are quantified in Fig 6F. **(D)** CAPRIN1-tethered ISR translation response for all reporters. Plotted as in Fig 6G for all reporters. For comparison, results of Fig 6G are replotted. **(E)** Average ISR translation response across all CAPRIN1-tethered reporters. Plotted are the average TE values across all reporters shown in (D). **(F)** CAPRIN1-tethered ISR protein response for all reporters. Plotted is the fold change in luciferase protein in untethered cells (green), tethered (+SGs) cells (pink), and tethered (–SGs) cells (red) during a time course of treatment with arsenite stress (500 µM sodium arsenite). Luciferase levels are reported relative to that of unstressed cells from the same biological replicate; otherwise as in (D). **(G)** Average ISR protein response across all CAPRIN1-tethered reporters. Plotted are the average luciferase values across all three reporters shown in (F). **(H)** CAPRIN1-tethered ISR RNA response for all reporters. Plotted is the fold change in reporter mRNA in untethered cells (green), tethered (+SGs) cells (pink), and tethered (–SGs) cells (red) during a time course of treatment with arsenite stress (500 µM sodium arsenite). RNA levels are reported relative to those of unstressed cells from the same biological replicate; otherwise as in (D). **(I)** Average ISR RNA response across all CAPRIN1-tethered reporters. Plotted are the average RNA fold changes across all reporters shown in (H).

